# Task-evoked deactivations: dissociation between BOLD fMRI and FDG

**DOI:** 10.64898/2026.05.14.725188

**Authors:** Tyler Blazey, John J Lee, Abraham Z Snyder, Marcus E Raichle, Hongyu An, Manu S Goyal, Andrei G Vlassenko

**Affiliations:** Mallinckrodt Institute of Radiology, Washington University School of Medicine, St. Louis, MO 63110; Neuroimaging Labs Research Center, Washington University School of Medicine, St. Louis, MO 63110; Department of Neurology, Washington University School of Medicine, St. Louis, MO 63110; Knight Alzheimer Disease Research Center, Washington University School of Medicine, St. Louis, MO 63108; Department of Neuroscience, Washington University School of Medicine, St. Louis, MO 63110; Department of Biomedical Engineering, Washington University in St. Louis, St. Louis, MO 63130; Department of Psychology & Brain Science, Washington University in St. Louis, St. Louis, MO 63130; Department of Electrical and Systems Engineering, Washington University in St. Louis, St. Louis, MO 63130

## Abstract

Task-evoked decreases in blood-oxygenation-level-dependent (BOLD) signals are a well-recognized phenomenon in functional magnetic resonance imaging (fMRI) studies. These deactivations are most prominent in the default mode network (DMN), a set of regions most active at rest. The metabolic basis of task-induced BOLD fMRI deactivations remains unclear. To address this question, we used PET/MRI to simultaneously measure BOLD fMRI and cerebral glucose consumption (CMRglc) during visuomotor and language tasks in 22 cognitively unimpaired older adults (15 female, 7 male). Task performance increased BOLD signals in task-relevant regions and decreased BOLD signals in the DMN. Positive BOLD responses generally coincided with increases in CMRglc. In contrast, CMRglc did not decrease in regions showing negative BOLD responses; instead, it typically increased. In particular, the posterior cingulate cortex showed significant CMRglc elevations in conjunction with negative BOLD responses. Whole-brain intensity normalization partially restored task-induced decreases in CMRglc, indicating that relative reductions appear in regions in which CMRglc increases are smaller than the global average. Overall, our results imply that BOLD fMRI deactivations can occur in conjunction with stable or even increased glucose consumption.

## Introduction

Since the beginning of functional neuroimaging, investigators have observed focal decreases in cerebral blood flow (CBF) during periods of imposed sensory stimulation or cognition (Frith et al., 1991). These findings gained prominence after Shulman et al. demonstrated that a range of tasks consistently reduced CBF in a set of regions including the precuneus/posterior cingulate (PCC), inferior parietal lobe, and medial prefrontal cortex (Shulman et al., 1997). Subsequent functional magnetic resonance imaging (fMRI) studies identified tasked-evoked decreases in blood-oxygenation-level-dependent (BOLD) signals within the same regions (McKiernan et al., 2003; Greicius and Menon, 2004). Together, these regions were later recognized as core components of the default mode network (DMN), a widely distributed constellation of regions exhibiting CBF and BOLD signal decreases during most goal-directed tasks (Raichle et al., 2001).

The metabolic and neuronal mechanisms underlying task-evoked decreases in CBF and BOLD signals remain an active area of investigation. One interpretation is that deactivations reflect reduced neuronal activity. Task-evoked decreases in local field potentials have been observed in both humans (Lachaux et al., 2008; Jerbi et al., 2010; Jung et al., 2010) and animals (Hayden et al., 2009; Popa et al., 2009), consistent with the correlation between the BOLD signal and local field potentials (Logothetis et al., 2001). However, the relationship between neuronal activity and local field potentials is complex (Buzsáki et al., 2012), and studies employing single unit recordings show both decreases and increases in association with BOLD deactivations (Hayden et al., 2009; Popa et al., 2009; Laurent et al., 2025).

If neuronal activity decreases during task-evoked deactivations, one would expect a corresponding decrease in cerebral glucose consumption (CMRglc), given its close relationship with neuronal activity (Sokoloff, 1999). [^18^F]fluorodeoxyglucose (FDG) PET has recently been used to investigate task-evoked deactivations in a series of studies by Andreas Hahn and colleagues (Hahn et al.,2016, 2018; Stiernman et al., 2021; Godbersen et al., 2023). These studies demonstrated that CMRglc changes in the PCC vary by task, increasing (Stiernman et al., 2021; Godbersen et al., 2023), decreasing (Hahn et al., 2016, 2018; Godbersen et al., 2023), or remaining unchanged (Hahn et al., 2016).

The studies by Hahn and coworkers employed the functional PET (fPET) technique, in which tracer is infused continuously throughout the scan rather than administered as a single bolus (Villien et al., 2014). The primary advantage of this approach is a substantial improvement in temporal resolution. With the traditional bolus method, only a single estimate of CMRglc can be obtained across the entire scan period. Consequently, measuring task-evoked responses typically requires separate rest and task scans separated by hours or days. In contrast, fPET enables quantification of CMRglc across multiple conditions within a single scan, with condition blocks as brief as a few minutes (Rischka et al., 2018). This eliminates between-session variability and enables detection of more transient metabolic changes.

However, fPET has disadvantages relative to the bolus-approach, most notably the need to model the “baseline” fPET signal in order to extract task-evoked changes (Villien et al., 2014; Hahn et al., 2016). Importantly, there is no universally accepted baseline model, whereas the choice of model can influence detected task-evoked changes (Coursey et al., 2024). With this in mind, we sought to determine whether: 1) task-evoked decreases in CMRglc are also observed with the traditional single bolus FDG PET technique, and 2) changes in CMRglc within the DMN are as task-dependent with the bolus method as they are with fPET.

## Materials and Methods

### Participants

Twenty-three cognitively unimpaired (CDR 0) older adults were recruited from the Knight Alzheimer Disease Research Center and the greater St. Louis community. We were unable to scan one participant due to size constraints, leaving us with a sample size of 22 individuals (7 male, 15 female) with a mean age of 72.6 years (SD 4.2; Range 65-78). Average height and weight were 164.7 cm (SD 8.2) and 85.1 kg (SD 22.3). All participants reported their ethnicity as Non-Hispanic or Latino. Among them, 20 self-identified their race as White, two as Black or African American, and one as Asian. Twenty participants identified as predominantly right-handed and two as predominantly left-handed. Written informed consent was obtained from all participants. All study procedures were performed according to the principles of the Declaration of Helsinki and received approval from the Washington University School of Medicine Institutional Review Board and Radioactive Drug Research Committee.

### Experimental Design

This study employed a within-subjects design, with each participant taking part in all three conditions: rest (eyes-open), a visuomotor task, and a word-stem completion task (WSCT; **Figure 1**). Each condition was administered on a separate day and took ~1 hour, with the task period occurring in the first 30 minutes of the scan. Condition order was varied across participants, although the rest condition was prioritized so that, except in one participant, it occurred during the first two visits. Fourteen participants performed the WSCT task before the visuomotor task, and 10 participants completed the rest condition before either task condition.

**Figure 1.**
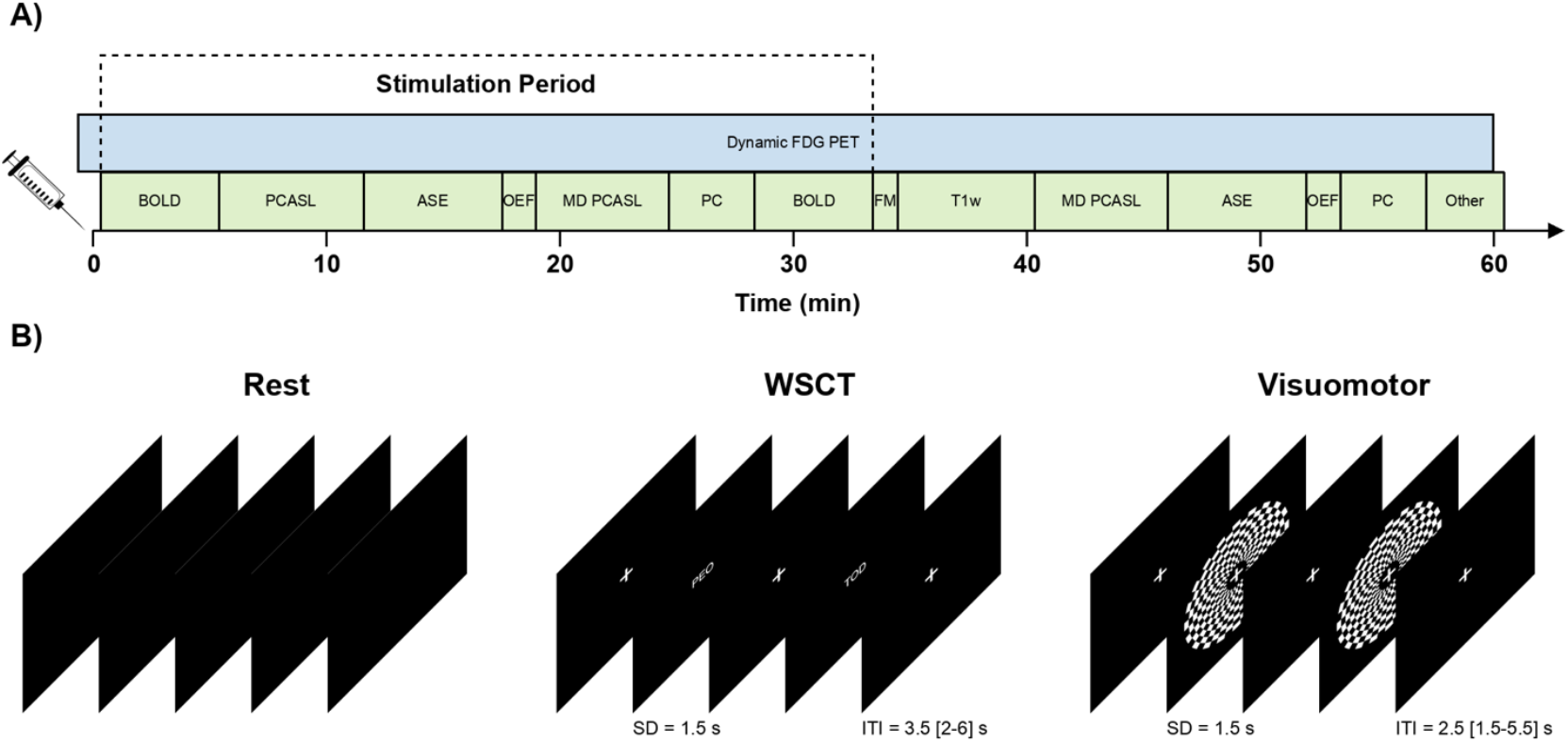
Experimental paradigm. A) Timing of stimulation, MRI scans, and PET scan relative to tracer injection. B) Example of the tasks participants performed during the stimulation period.

Participants were instructed to fast for six hours before each visit. Upon arrival, an IV was placed in each forearm: one to inject a nominal dose 185 MBq of FDG, and the other to obtain up to three venous blood samples during the final 30 minutes of the PET scan. Fasting plasma glucose was also measured to ensure that the participant was not hyperglycemic. The mean glucose level was 100.6 mg/dL (SD 15.7, range 79.0 – 164.0). Before entering the MRI room, the participant had the task procedure explained to them and had the opportunity to ask questions. They were then moved into the scanner room, where they were instructed on the use of the response box (Current Designs, HHSC-1×4-L). Blood oxygen saturation (mean 96.2%, SD 1.6%) was measured using an MR-compatible pulse oximeter (Phillips, Expression MR400).

Once they were in the scanner, participants were told to minimize movement and to keep their eyes open. During task visits, they were also shown task instructions and completed a brief practice session. The experiment proceeded only when participants successfully completed the practice trials (i.e., pressed the response button within the trial interval) and verbally confirmed their understanding of the task. The task began, on average, 62 seconds (SD 17, range 36 – 107) from the start of the FDG injection.

### Cognitive Tasks

During the visuomotor task, participants were told to press a button with their right index finger every time they saw a rotating circular checkerboard (http://cognitiveatlas.org/task/id/trm_4c898f8f297ac/). The checkerboard was shown for 1.5 seconds during each trial, followed by a jittered intertrial interval (1.5 - 5.5 s) where only a white fixation cross was shown. The distribution of 485 intertrial intervals was optimized so that it approximately followed an exponential distribution with a mean of 2.5 seconds. A ten-second fixation period was also added to the start and end of each BOLD run to facilitate estimation of the baseline signal.

The WSCT is a more demanding language processing task (https://www.cognitiveatlas.org/task/id/trm_4c8a858da803d/). Participants are shown a three-letter word stem (e.g., HOU) and asked to think of a word that completes it (e.g., HOUSE). Participants were specifically instructed to press a button with their right index finger as soon as they thought of the word, and not to speak the word out loud. We chose not to use an overt language task to minimize participant movement (Barch et al., 1999) and other imaging artifacts associated with speaking (Birn et al., 1998; Scholkmann et al., 2013).

The timing for the word-stem completion task (WSCT) was similar to the visuomotor task, with each word-stem being shown for 1.5 seconds and then followed by a jittered intertrial interval (mean 3.5 s, range 2 – 6.5 s). The entire task included 388 trials, each one with a unique English word stem. Each word stem was included in a list of the 5,000 most frequent English words (drawn from the Corpus of Contemporary American English, a database of over one billion words from over 485,202 texts, https://www.english-corpora.org/coca/). The median stem frequency was 328 appearances per million words, with frequences ranging from 12 (UNB) to 15,564 (THA) per million.

Both tasks were programmed using PsychoPy 2023.1.3 (Peirce et al., 2019) and are available to download at https://github.com/tblazey/py_task.

### MRI Acquisition

All images were acquired using a Siemens Biograph 3T PET/MR equipped with a 32-channel head coil. During the task block, six different MR sequences were acquired in order. The first, which also repeated at the end of the task-period, was a 2D echo-planar imaging (EPI) sequence for measuring task-evoked BOLD activity. This sequence consisted of 150 volumes acquired with a repetition time (TR) of 2 s, a flip angle of 70 degrees, an echo time (TE) of 28 ms, an echo spacing of 0.6 ms, a 64 × 64 acquisition matrix with 32 slices, and a 4 mm isotropic voxel size. The phase encoding direction was anterior-to-posterior for the first run and posterior-to-anterior for the second run.

The first BOLD run was followed by a single-delay 2D pseudo continuous arterial spin labeling (pCASL) sequence for regional cerebral blood flow (Dai et al., 2008), and an asymmetric spin echo (ASE) sequence for regional oxygen extraction fraction (OEF; An and Lin, 2003). Neither sequence is reported here as they did not show robust changes in expected task-activated regions. Global oxygen extraction fraction (OEF) was then measured using T_2_-Relaxation-Under-Spin-Tagging (TRUST; Lu and Ge, 2008). A single 5 mm slice positioned across the posterior aspect of the superior sagittal sinus was acquired with a 3 s TR, 4.2 ms TE, inversion time (TI) of 1.02 s, 10 ms inter-echo spacing, 90° flip angle, GRAPPA acceleration factor of 3, and a 64 × 64 acquisition matrix with 11.5 mm^2^ in-plane resolution. We acquired twelve label–control pairs spanning four effective TEs (0.44, 40, 80, and 160 ms), with each TE measured in triplicate.

Next, we acquired a multi-delay 3D pCASL, which like the 2D pCASL, will not be included because it did not reliably reveal task-related increases in CBF. This was followed by 3D phase contrast (PC) MRI for global CBF (Wymer et al., 2020). This sequence included 3 orthogonal 75 cm/s velocity encodings acquired with a TR of 40.65 ms, a TE of 5.60 ms, a flip angle of 10°, a GRAPPA acceleration factor 2, 384 × 512 × 128 acquisition grid, and 0.5 × 0.5 × 0.9 mm voxel size. The acquisition slab was centered around the upper cervical spine to ensure coverage of the internal carotid (ICA) and vertebral arteries.

A gradient-echo field map was acquired immediately after the task period with a 488 ms TR, 5.19- and 7.65 ms TEs, 60° flip angle, 64 × 64 acquisition matrix, 32 slices, and 4-mm isotropic voxels. A high-resolution T_1_-weighted (T_1_w) image of the head was then obtained using a magnetization-prepared rapid gradient-echo (MPRAGE) sequence with a 2.4-s TR, 2.21 ms TE, 8° flip angle, 0.9-mm isotropic voxels, 2× GRAPPA acceleration, 288 × 272 acquisition matrix, and 192 slices.

After the field map and MPRAGE we repeated the multi-delay pCASL, ASE, TRUST, and PC scans to estimate resting metabolism during the same visit. The final portion of the MR protocol varied between visits. During the visuomotor visit, a 3D time-of-flight (TOF) magnetic resonance angiogram (MRA) was acquired to segment the internal carotid artery for PET image-derived arterial input functions. The sequence used a 23 ms TR, 3.6 ms TE, 18° flip angle, and 168 slices at 0.6-mm thickness. The initial in-plane resolution was 0.6 mm isotropic (384 × 288 acquisition matrix) which was then up sampled by a factor of two to yield a 0.3-mm isotropic resolution. A high resolution 3D T_2_-weighted images (3 s TR, 407 ms TE, 120° flip angle, 256 × 256 acquisition matrix with 224 slices, 1 × 1 × 0.9 mm voxels) during the rest visit.

### MRI Image Analysis

Processing began with the sMRIPrep 0.16 (RRID:SCR_016216; Esteban et al., 2019), a standardized reconstruction pipeline based upon NiPype 1.8.6 (RRID:SCR_002502; Gorgolewski et al., 2011). Each T_1_w image was corrected for intensity non-uniformity (INU) with N4BiasFieldCorrection (Avants et al., 2011), distributed with ANTs 2.5.3 (RRID:SCR_004757; Avants et al., 2008). An anatomical T_1_w-reference map was computed after registration of all 3 visits T_1_w images (after INU-correction) using mri_robust_template (Reuter et al., 2010) in FreeSurfer 7.3.2 (RRID:SCR_001847; Fischl, 2012). The T_1_w-reference was then skull-stripped with a Nipype implementation of the antsBrainExtraction.sh workflow (from ANTs), using OASIS30ANTs as target template. Brain tissue segmentation of cerebrospinal fluid (CSF), white-matter (WM) and gray-matter (GM) was performed on the brain-extracted T_1_w using fast (Zhang et al., 2001) in FSL (RRID:SCR_002823; Jenkinson et al., 2012). Brain surfaces were reconstructed using recon-all (Dale et al., 1999), and the brain mask estimated previously was refined with a custom variation of the method to reconcile ANTs-derived and FreeSurfer-derived segmentations of the cortical gray-matter of Mindboggle (RRID:SCR_002438; Klein et al., 2017). A T_2_-weighted image was used to improve pial surface refinement. Volume-based spatial normalization to standard space (RRID:SCR_002823; TemplateFlow ID: MNI152NLin6Asym) was performed through nonlinear registration with antsRegistration (ANTs 2.5.3), using brain-extracted versions of both T_1_w reference and the T_1_w template.

BOLD image processing was performed using FEAT (FMRI Expert Analysis Tool) in FSL 6.0.6.1 (RRID:SCR_002823; (Jenkinson et al., 2012). Preprocessing included motion correction with MCFLIRT (Jenkinson et al., 2002), distortion correction with PRELUDE (Jenkinson, 2003) and FUGUE (Jenkinson et al., 2012), brain masking using BET (Smith, 2002), spatial smoothing with a 5 mm FWHM Gaussian kernel, grand-mean intensity normalization the entire 4D dataset, and high-pass temporal filtering (sigma 50 s). Four runs were excluded for excessive motion, defined as mean absolute displacement > 1 mm or mean relative displacement > 0.4 mm. An additional five runs were excluded because the number of missed responses exceeded 2 SD of the mean (WSCT: mean = 3.6, SD = 6.3; Visuomotor: mean = 6.3, SD = 11.7).

First-level (i.e, timeseries) analysis was performed using FILM (Woolrich et al., 2001) with correction for local temporal autocorrelation. The design matrix included a task regressor, modeled as each 1.5 s event convolved with a double-gamma hemodynamic response function, its temporal derivative, and the six motion parameters estimated by MCFLIRT. After fitting, the image of voxelwise regression coefficients for the task effect was resampled to standard space. For each run, the temporal average BOLD image was rigidly aligned to the sMRIPrep-generated T1w template using FLIRT (Jenkinson and Smith, 2001) using a gray/white boundary-based cost function (Greve and Fischl, 2009). This linear transformation was then applied with the nonlinear transformation estimated by sMRIPrep to bring each parameter estimate into MNI152NLin6Asym space without changing the voxel size. Region of interest (ROI) averages were then computed for each run’s task effect image. The ROI atlas comprised 200 cortical gray matter parcels (Schaefer et al., 2018) (Figure S1) and 16 subcortical gray matter regions (Tian et al., 2020) defined by resting state connectivity. The region set also included 32 cerebellar gray regions from a hierarchical cerebellar atlas (Nettekoven et al., 2024), created by fusing segmentations derived from resting state and task MRI data.

Whole-brain OEF was computed from the TRUST MRI data using methods adapted from the work of Lu and colleagues (Lu and Ge, 2008; Lu et al., 2012; Gou et al., 2024) and implemented in a custom numerical Python (RRID:SCR_008633; Harris et al., 2020; Virtanen et al., 2020) script (https://github.com/tblazey/trust). First, we first calculated the difference between control and label images for each effective TE pair. The three pairs at the minimum TE (0.44 ms) were averaged to generate an ROI in the superior sagittal sinus, defined as the four highest-intensity voxels within a 3 × 3 grid centered on the voxel with the greatest signal. All ROIs were manually reviewed to confirm their location within the superior sagittal sinus. A negative exponential model was fit to the ROI-averaged data for each difference image to estimate the T_2_ relaxation time of blood (T_2b_) (Lu and Ge, 2008). For fitting purposes, the T_1_ relaxation time of blood was assumed to be 1.624 s. T_2b_ was then converted to venous oxygenation (Y_v_) using a polynomial relationship between T_2b_, hematocrit and Y_v_ (Lu et al., 2012). OEF was computed as the percent difference between Y_v._ and Y_a_, where Y_a_ is the arterial oxygen saturation measured with a pulse oximeter (Jiang et al., 2021).

To facilitate computation of whole-brain average CBF, participant-specific PC MRI template images were created by rigidly registering each visit’s magnitude PC image. Each participant’s PC template was then rigidly aligned to their T_1_-weighted template. The sum-of-squares magnitude-difference image for each visit was transformed into atlas space by combining these two transformations with the T_1_w-to-atlas warp generated by sMRIPrep. An average magnitude-difference image was then computed in atlas space. This image was used to manually identify a slice inferior to the V3 segment of the vertebral artery where both sides of the internal carotid and vertebral artery were visible. For each artery, an ROI was created by manually identifying a center voxel and fitting a circle to the surrounding voxels. These atlas-space ROIs were transformed to native space, dilated by a single voxel, and used to identify a visit-specific peak voxel around which to fit an updated circular ROI. This last step was done to ensure that misregistration between visits did not cause ROIs to include voxels outside of the target vessel. We also manually inspected each ROI to confirm its placement.

Blood flow (mL/min) was computed by multiplying the average sum-of-squares phase difference within each arterial ROI by the ROI area and the velocity-encoding factor (75 cm/s). Flows from all ROIs were then summed and divided by total brain volume to yield whole-brain CBF. Total brain volume was estimated using the ‘BrainSegNotVent’ value generated by FreeSurfer, which includes the cerebellum and cerebrum but excludes the ventricles and brain stem.

Whole-brain cerebral metabolic rate of oxygen (CMRO_2_) was calculated as:

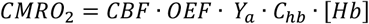

where CBF and OEF were estimated using PC and TRUST MRI, respectively, *[HB]* is the total hemoglobin concentration, and *C*_*hb*_ is the oxygen carrying capacity of hemoglobin (59.8 µmol O_2_/gram hemoglobin; Jiang and Lu, 2022).

To facilitate extraction of the PET arterial input function, the Segment 3D tool in ITK-SNAP (RRID:SCR_002010; Yushkevich et al., 2006) was used to semi-automatically segment the internal carotid artery from the MRA. The magnitude TOF MRA was manually thresholded to remove low-intensity voxels unlikely to contain large vessels. A contour evolution algorithm, with default settings and seed points placed manually along the ICA, was then used to segment both internal carotids. The number of iterations was adjusted empirically for each case to ensure the entire visible ICA was included in the final segmentation. The ICA mask was then transformed to atlas space via linear registration of the TOF MRA to the participant’s T_1_w template.

### PET Acquisition and Analysis

A 65-minute dynamic PET scan starting on average 37 s (SD 10) before the injection of 195.0 MBq (SD 12.5) of [^18^F]FDG was acquired at each visit. The scanner was started early to ensure that the bolus arrival was captured. Emission images were reconstructed using on-scanner tools, which included correction for scatter, randoms, dead time, and attenuation. Attenuation maps were obtained using a Dixon-based MRI sequence (Dixon, 1984) provided by Siemens. Image reconstruction was performed using a 3D ordered-subset expectation maximization (OSEM) algorithm using variable frame durations (24 × 5 s, 9 × 20 s, 10 × 60 s, and 10 × 300 s).

The dynamic PET data was motion corrected by first spatially smoothing each 4D PET frame with a 5 mm FWHM Gaussian kernel and then computing the spatial correlation between each frame and the final frame, which served as a reference. Frames with a Pearson correlation coefficient > 0.5 were rigidly aligned to the reference using FLIRT (Jenkinson and Smith, 2001); those below this threshold were not realigned. Across all scans, an average of 88.9% (SD = 2.8) of frames with counts were realigned. A participant specific FDG template was then created by skull-stripping each visits reference frame with SynthStrip (Hoopes et al., 2022) then aligning them with mri_robust_template (Reuter et al., 2010). A rigid body registration was computed between each participant’s FDG template and their T_1_w template image using FLIRT. The unsmoothed 4D PET data from each visit was then transformed to MNI152NLin6Asym space without modifying the voxel size by combining motion-correction, rigid transformations, and nonlinear warp in one step.

Average time-activity curves were then computed using the same 248 ROI atlas used for the BOLD data and an atlas-based whole-brain mask.

### Image-derived Arterial Input Functions

Our primary method for estimating arterial input functions was based on the method used by the EMATA toolbox (https://github.com/FairUnipd/EMATA; De Francisci et al., 2024). First, we extracted all the time-activity curves for all the voxels within the TOF MRA–defined ICA mask. To exclude voxels unlikely to be arterial, we first removed voxels whose temporal peak deviated by more than 2.5 seconds from the median across all masked voxels. Each voxel was then summarized using eight features: peak amplitude, mean of the last four points, rising slope, ending slope, area before the peak, area after the peak, and standard deviation. Principal component analysis reduced these features to four dimensions, which were then classified into two groups with K-means clustering (RRID:SCR_002577; Pedregosa et al., 2011). We averaged the 100 voxels with the highest individual peaks from the cluster whose center had the higher peak amplitude to create the final arterial input function.

To confirm the robustness of our results, we performed a *post hoc* supplemental analysis using imaged-derived arterial input functions extracted using a method based on that of Sari et al. (**Figure S2**; Sari et al., 2017). The main steps of this method are: 1) creating an ICA mask and 2) partial volume correction of the PET signal within the ICA mask using a single-target approach. We introduced two modifications. First, we manually refined the ICA mask generated for the EMATA-based input functions to ensure proper alignment with the arterial signal in the PET image. This step was needed because the MR angiogram was acquired only during the visuomotor visit, making registration to the rest and WSCT visits more difficult. Second, we applied HYPR denoising (Floberg et al., 2012) before partial volume correction. Without denoising, the single-target correction often produced negative values due to noisy voxels outside the arterial mask. Denoising was used only for input function extraction; regional data used for kinetic modeling were not denoised.

### Kinetic Modeling

The extracted input functions were then fit to a reversible 2-compartmnet model:

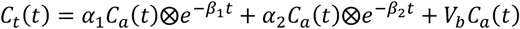

Where *α*_*1*_, *α*_*2*_, *β*_*1*_, and *β*_*2*_ are products of the fractional blood volume, *V*_*b*_, and the traditional rate constants (*K*_*1*_, *k*_*2*_, *k*_*3)*_, and *k*_*3*_) (Rizzo et al., 2013), *C*_*a*_ is the imaged-derived arterial input function, and *C*_*t*_ is the PET time-activity curve. Although *C*_*a*_ is often separated into whole-blood and plasma components, using a population-based corrected factor (Phelps et al., 1979) doing so had a minimal impact on task-evoked differences. Nonlinear least squares was used to estimate the five free model parameters (*α*_*1*_, *α*_*2*_, *β*_*1*_, *β*_*2*_, and *V*_*b*_) for each ROI. Uniform weights were applied due to uncertainty in the appropriate weighting scheme (Yaqub et al., 2006; Dai et al., 2011). Fitting was performed using optimization tools in SciPy (RRID:SCR_008058; Virtanen et al., 2020). CMRglc was calculated using the traditional formula: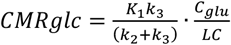, where *C*_*glu*_ is the plasma glucose concentration and LC is the lumped constant, which was set to 0.81(Wu et al., 2003).

The whole-brain oxygen-to-glucose index (OGI) was calculated by dividing the MR-based whole-brain CMRO_2_ by the CMRglc derived from the whole-brain average time-activity curve. We also computed standardized uptake value ratio (SUVR) for each ROI using the whole brain as a reference region to examine regional differences without the impact of any global changes. SUVR values were calculated by integrating the frames acquired 40 minutes past the start of scanning.

### Statistical Analyses

All statistical analyses were conducted using hierarchical linear models using brms 2.22.0 (RRID:SCR_023862; (Bürkner, 2017), an R 4.4.1(RRID:SCR_001905; (R Core Team, 2024) front end for STAN (RRID:SCR_018459; (Stan Development Team, 2024), a package for Bayesian inference with Hamilton Markov chain Monte Carlo. For whole-brain average parameters, the models was:

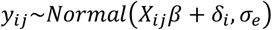

where *y*_*ij*_ is the data for participant *i* at timepoint *j, X*_*ij*_ is a 1 × 3 row vector including the intercept and task regressors, *β* is a 3 × 1 column vector of coefficients, *δ*_*i*_ is the participant-specific mean offset, and *σ*_*e*_ is the residual standard deviation. For measurements repeated within session (OEF, CBF, and CMRO_2_), a visit level intercept was added to the model. Generally, broad priors were chosen according to the default recommendations in brms. The intercept had a Student-T prior with a mean equal to the sample mean of *y*, a scale equal to the medial absolute standard deviation of *y*, and 3 degrees of freedom. The participant and visit level intercepts were given Normal priors with zero means and standard deviations *σ*_*δ*_ and *σ*_*η*_, respectively. The prior for the three standard deviation parameters was a half Student-T with zero mean and the same scale and degrees of freedom as the intercept prior. The only prior we modified from the default recommendations was for the task regressors, which was changed from a uniform distribution to a Normal distribution with mean 0 and standard deviation 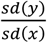.

Models for the regional data followed the approach pioneered by Chen et al., in which all regions are fit together within a single model (Chen et al., 2019). By treating each regional effect as a sample from a shared population-level distribution, this approach effectively mitigates the multiple comparisons problem. The specific model was:

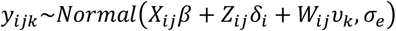

Where *y*_*ij*,_ is the data from ROI *k* for participant *i* at timepoint *j, Z*_*ij*_ and *W*_*ij*_ are row vectors containing participant and region-level regressors, and *δ*_*i*_ and *υ*, are column vectors of participant and region-level coefficients. The regressors were the same at all three levels for both BOLD and CMRglc. The CMRglc regressors included an intercept and a term for each task, while BOLD only used an intercept and a WSCT term because there was no resting scan. We also ran an analysis where a run regressor and a WSCT × run interaction was added to the BOLD model to determine if there was a difference in task-evoked activity between the beginning and end of the task period. For SUVR, we used the same population and region-level regressors as for CMRglc. However, because each SUVR image is whole-brain normalized, the participant-level terms only included an intercept nested within region.

The regional models used the same prior strategy for the population-level coefficients and the residual standard deviation as whole-brain brain models. However, because the higher levels included both intercepts and slopes, we gave their coefficients a multivariate normal prior:

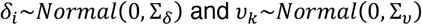

where Σ_*δ*_ and Σ_*υ*_ are the covariance matrices for the participant and region-level effects, respectively. For setting priors, each covariance matrix was decomposed into a vector of standard deviations and a correlation matrix (Barnard et al., 2000). The standard deviations were given the same half Student-T prior as the previous models. The prior for the correlation matrix was a LKJ correlation distribution (Lewandowski et al., 2009) with a shape of 1, which is effectively a uniform prior over all correlation matrices.

The primary parameters of interest were the population-level task coefficients in the whole-brain models. For the regional models, we focused on the sum of the population-level and region-level task coefficients for each ROI. This formulation captures the quantitative change in each region, with the population-level terms representing the effect common to all regions and the region-level terms providing region-specific offsets. Our primary outcome measure was whether the 95% highest density interval (HDI) for a comparison excluded zero (Kay, 2024). We also evaluated two complementary metrics: the probability of direction, defined as the fraction of the posterior distribution with the same sign as its median (Makowski et al., 2019), and the probability that the percent change exceeded −5%.

Models were run with 4 parallel chains, each with 10,000 iterations, half of which were warm-up iterations. The final posteriors therefore consisted of 20,000 samples. To assess sample quality, we used both the effective sample size, 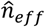, and the split scale reduction factor, 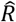 (Gelman et al.,2013). 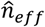, which is an estimate of the number of independent samples in each parameter’s posterior distribution, was always above 3,971 samples for all the task effects. At convergence, *R*, which is a measurement of within chain variance to the pooled between chain variance, should be equal to 1. Across all models, *R* was never greater than 1.00 for all task effects. We also performed posterior predictive checks to assess whether the model provided a reasonable representation of the data (Gelman et al., 2013). For each posterior sample, we simulated CMRglc values for a new subject - drawn from a multivariate normal with zero mean and covariance matrix equal to Σ_*δ*_ - across four ROIs during all three conditions (**Figure 3**).

All the data and code necessary to reproduce the figures and tables in this report can be found at http://doi.org/10.5281/zenodo.20073537.

## Results

### Whole brain metabolism

**Table 1** provides quantitative estimates of task-evoked changes in cerebral metabolism for the entire brain. We observed no robust changes in any whole brain parameter between rest and either the visuomotor or WSCT condition. However, on average whole brain CMRglc increased during both tasks (visuomotor: 2.6% [-7.8, 13.6], WSCT: 8.0% [-2.8, 19.3]), with a trend-level change during the WSCT condition (*P*(Δ > 0) = 0.93). There were also trend level decreases in whole brain OGI for both tasks and in OEF during the visuomotor task. Excluding CMRglc, average whole brain resting parameter values were lower compared to previous estimates in the literature (Fan et al., 2016; Blazey et al., 2018; Otto Henriksen et al., 2018; Henriksen et al., 2021), indicating a bias in our MR-based estimates.

**Table 1.**
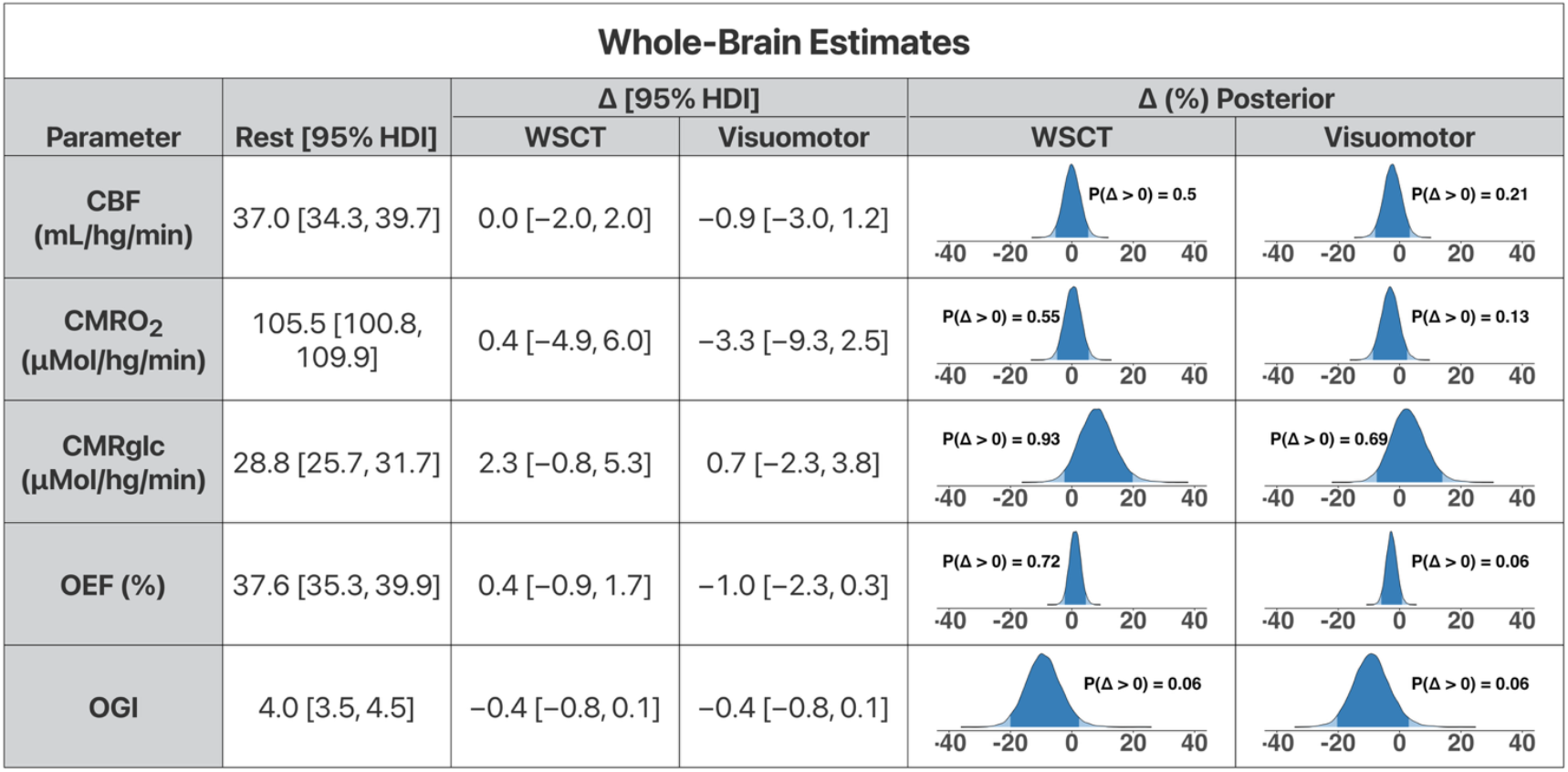
Quantitative whole-brain parameter estimates. The first column is the resting mean and its 95% highest density interval (HDI) for cerebral blood flow (CBF), oxygen consumption (CMRO_2_), glucose consumption (CMRglc), oxygen extraction (OEF), and the oxygen-to-glucose index (OGI), The next two columns contain the average difference between rest and task. Final two columns are the posterior distribution of the % difference between each task and rest.

### Regional changes in CMRglc and BOLD

Consistent with the trend level increase in whole brain CMRglc, every gray-matter region examined showed greater CMRglc during the WSCT condition than during rest (**Figure 2**). The 95% HDI excluded zero in 45 of 248 regions, including areas, such as the visual cortex and left prefrontal cortex, where BOLD signals increased (**Figures 2, 3**). However, CMRglc also increased in the right lateral prefrontal cortex and bilateral posterior cingulate, regions where BOLD signals were unchanged or even decreased. To quantify the confidence that CMRglc did not decrease, we computed the probability that the percent change in CMRglc was greater than −5% (**Figure 4**). For most regions (188/248), including in parts of the DMN, this probability was greater than 95%. Exceptions included the medial temporal lobe, inferior parietal lobe, and medial prefrontal cortex, where the probability of a change greater than −5% usually exceeded 80% (55/60 regions). Widespread increases were also observed with an alternative imaged-derived arterial input function method, indicating that this result was not overly sensitive to our image processing approach (**Figure S2**).

**Figure 2.**
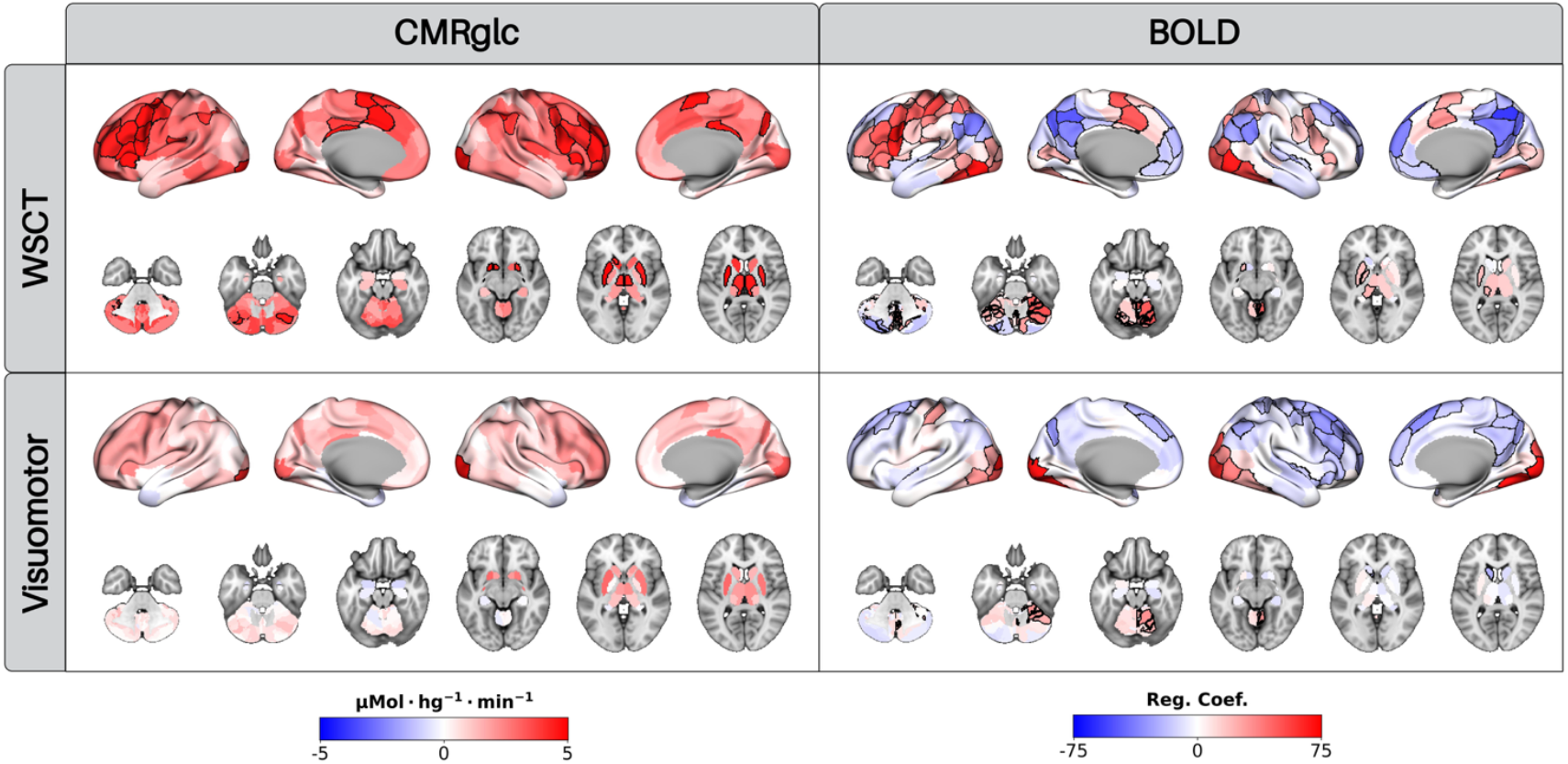
Task-evoked deactivations are present in fMRI BOLD but not CMRglc. The columns represent modalities (CMRglc and BOLD), while the rows correspond to tasks (WCST and visuomotor). Regions in which the 95% highest density interval (HDI) for the difference between rest and task does not overlap zero are outlined in black. Both tasks produced global increases in CMRglc, although our 95% HDI threshold was only met in a subset of regions. BOLD signal increases were more focal and varied according to task. Although no decreases CMRglc were observed for either task, there were multiple regions where we can conclude with confidence that was a decrease in BOLD signal (0 ∉ 95% HDI).

**Figure 3.**
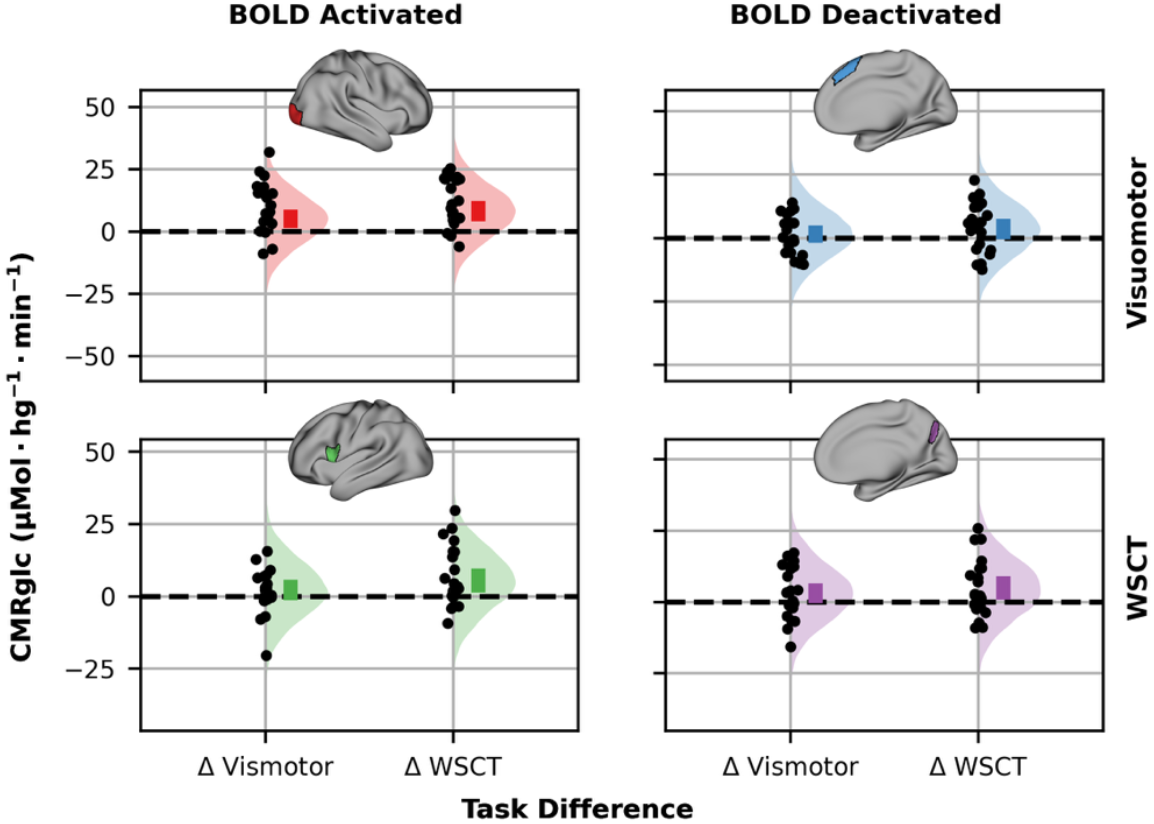
Difference in cerebral metabolism between rest and task for four regions of interest. Regions were selected based on regions in which BOLD signals showed task-related activations or deactivations (**Figure 2**). Each dot is the estimated difference from an individual participant. Colored bars are the 95% highest density interval (HDI) of the mean; the density plot is a posterior predictive distribution. Note that although there were increases in CMRglc during the task at the participant-level, decreases were not found at the group-level.

**Figure 4.**
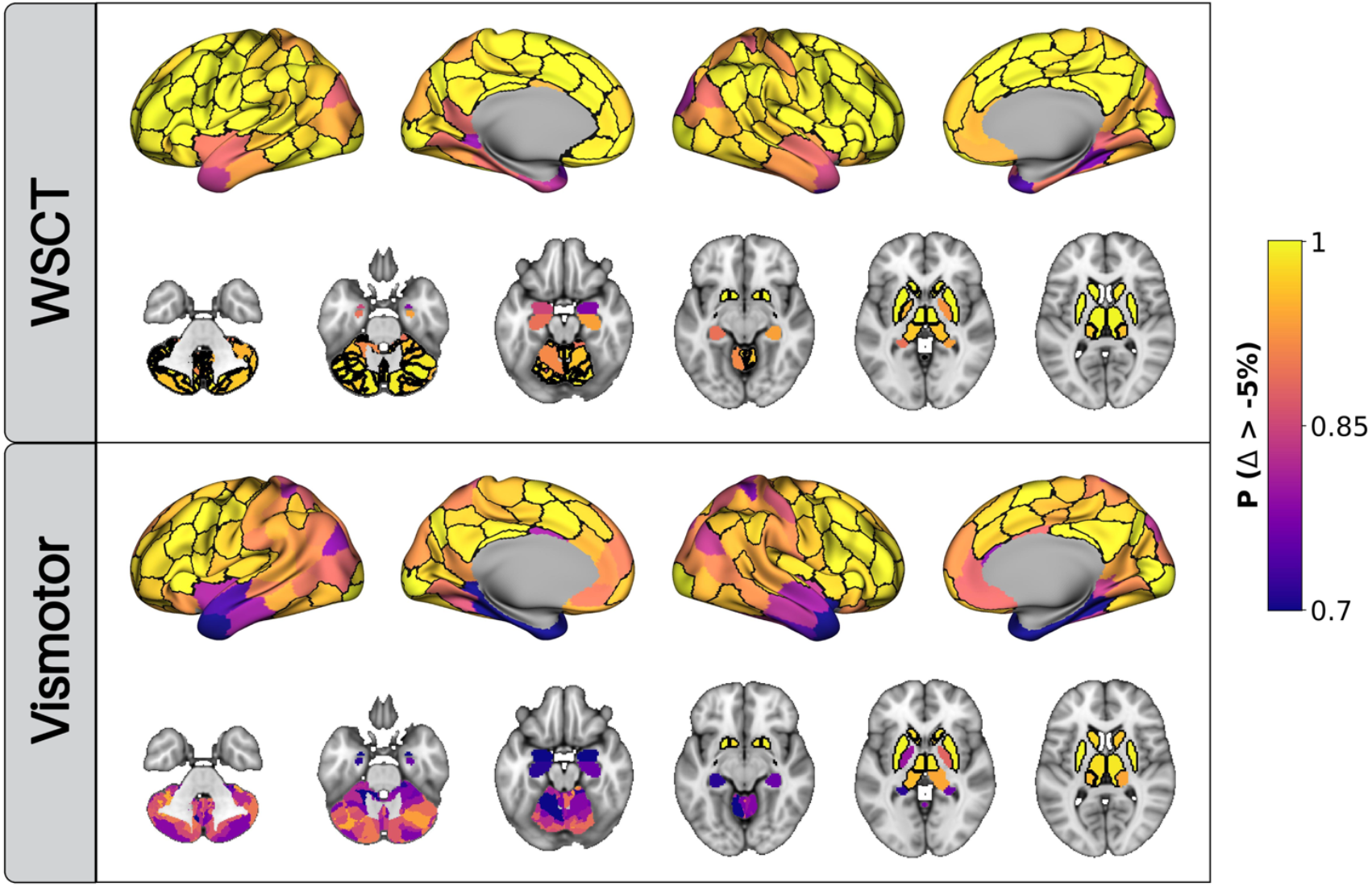
Large task-evoked decreases in CMRglc are unlikely. Maps display the probability that task-evoked change, Δ, is greater than −5% of rest. Regions where this probability is greater than 0.95 are outlined in black (WSCT:188/248, visuomotor:113/248).

The visuomotor task also increased CMRglc across the brain, although the visual cortex was the only region where the 95% HDI excluded zero (**Figure 2, 3**). BOLD signals in the visual cortex also increased during the visuomotor task, although the area of activation was broader than that of CMRglc. Surprisingly, the task-evoked decreases in the BOLD signal included both lateral and medial portions of the dorsal prefrontal cortex in addition to the expected DMN regions. Although, on average, CMRglc increased in regions where BOLD decreased, we could not rule-out a null effect (0 ∈ 95% HDI). However, large decreases in regional CMRglc were unlikely. The probability that the change in CMRglc exceeded −5% was greater than 0.95 in 113 out of 248 regions (**Figure 4**). In the remaining 135 regions, the probability of a change exceeding −5% was above 0.80 in 104 regions.

### Spatial overlap between changes in BOLD and CMRglc

We next computed conjunction maps to directly assess the spatial overlap between task-evoked changes in BOLD and CMRglc (**Figure 5**). The WSCT condition increased both BOLD signals and CMRglc throughout the prefrontal cortex, anterior cingulate, and visual cortex (*P*(Δ > 0) > 0.95; 46 regions). Using a 0.95 confidence level, ten ROIs belonging to the default and control networks showed both increased CMRglc and decreased BOLD during the WSCT. However, the number of regions in this set is highly dependent on the confidence level, as BOLD declined in many regions where the probability of an increase in CMRglc did not reach 0.95 (**Figure 5**). For example, in four parcels of the right prefrontal cortex, the probability of a BOLD decrease exceeded 0.95, while the probability of a CMRglc increase ranged from 0.84 to 0.93.

**Figure 5.**
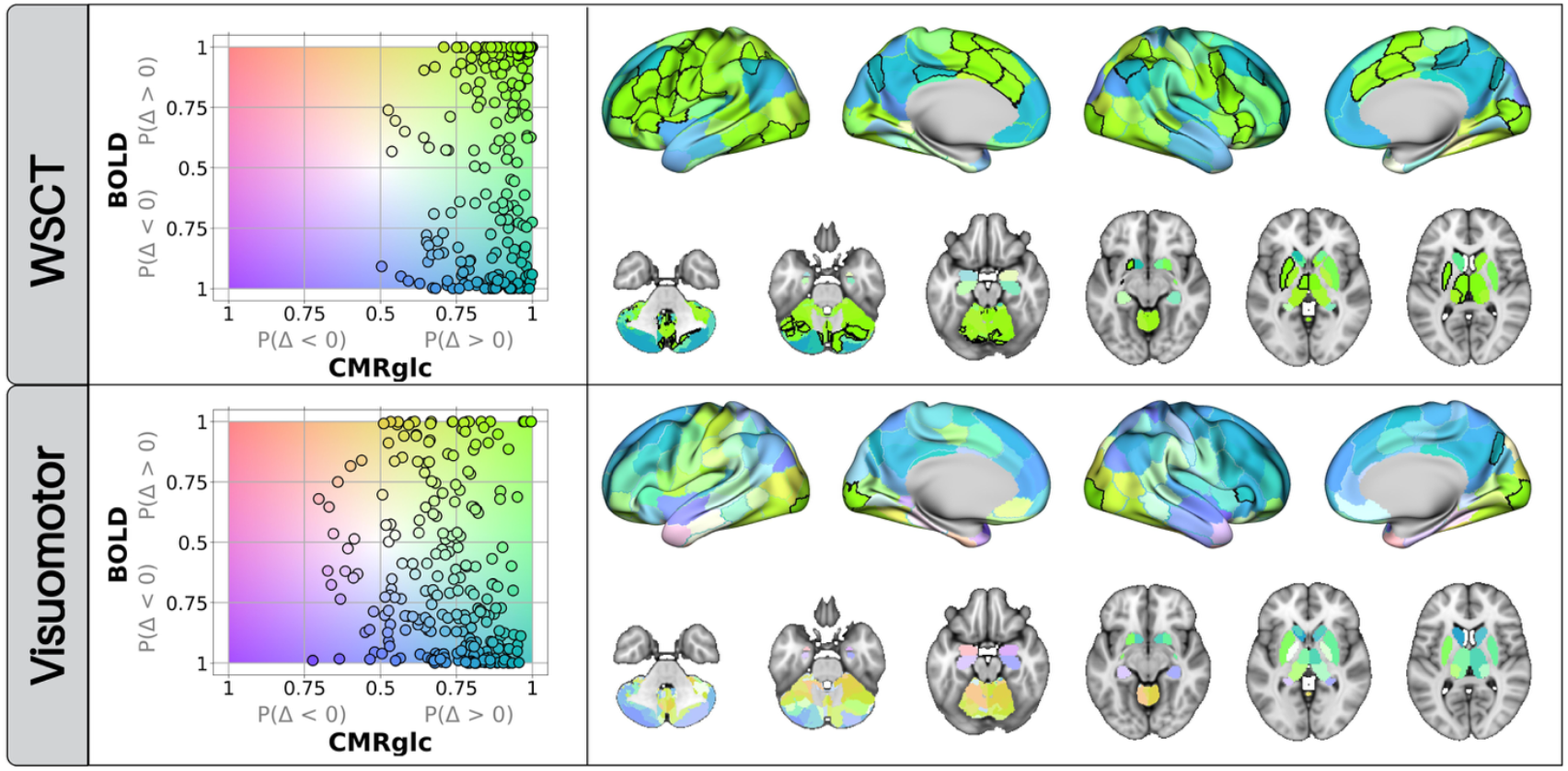
Spatial overlap between changes in CMRglc and BOLD signals. The column on the left shows the probability of a change for each modality plotted against each other. Each point is a region. Right hand column shows the same data plotted on the brain. All data is colored according to the 2D-colormap shown behind the scatterplot. For the WSCT condition, most regions occupy the right side of the color space, indicating a CMRglc increase along with an increase (green) or decrease (blue) in BOLD signal. There was more variably in the visuomotor task, with several regions showing evidence for only a decrease (light purple) or increase (yellow) in BOLD signal. Regions in which the probability of a decrease or increase exceeded 0.95 are outlined in black.

The conjunction maps for the visuomotor task were far sparser than the WSCT maps. Only four regions in the visual cortex showed elevated CMRglc and BOLD using a confidence threshold of 0.95. Similarly, only two parcels, one in the right precuneus and the other in the right insula, exhibited decreased BOLD along with increased CMRglc. As in the WSCT, there were many regions with a high probability of a decrease in BOLD signal and a moderate probability (*P*(Δ > 0) > 0.75) of an increase in CMRglc. Conversely, more regions showed BOLD signal increases (e.g., cerebellum) or decreases (e.g., medial temporal lobe) with little evidence of CMRglc change (*P*(Δ > 0) ~ 0.5).

### Relative decreases in glucose consumption

To account for the brain-wide increase in CMRglc during both tasks, we computed whole-brain normalized SUVR images of FDG uptake (**Figure 6**). Normalization revealed several regions where glucose metabolism decreased relative to the whole brain during both tasks (0 ∉ 95% HDI; WSCT = 56, visuomotor = 36). Although normalization increased the correspondence with fMRI, discrepancies between FDG and BOLD signals remained. In the right lateral prefrontal portion of the control network, SUVR increased during both tasks while BOLD signals were stable or decreased.

**Figure 6.**
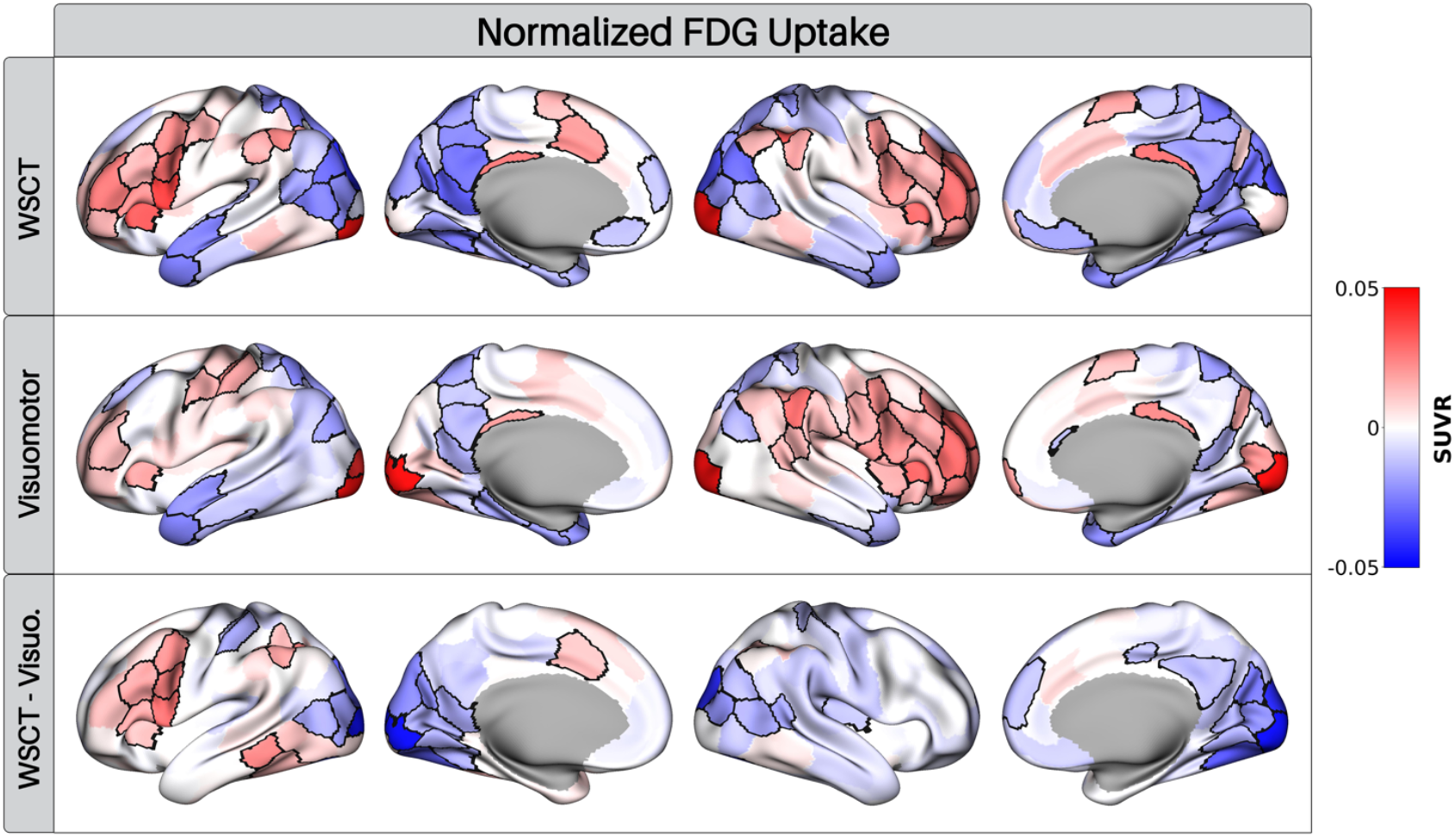
Whole brain normalized changes in task-evoked glucose consumption. Top two rows are the difference between rest and task. In addition to increases seen in **Figure 2**, normalization reveals several regions where glucose consumption decreased relative to the rest of the brain (e.g., precuneus). Subtraction of the visuomotor task from the WSCT eliminated the relative increase in right lateral prefrontal cortex glucose consumption seen in both tasks. Regions in which the 95% HDI does not include zero are outlined in black.

### Impact of prolonged task-performance on BOLD fMRI

The extended duration of our task-period raises the possibility of a change in BOLD signals or metabolism over the course of the task-period (Mintun et al., 2002; Vlassenko et al., 2006). Although time-varying CMRglc cannot be robustly estimated with a bolus injection, we acquired BOLD fMRI runs at both the beginning and end of the task period. The only region exhibiting a clear difference between the tasks is the right visual cortex, in which BOLD signals during the second run of the WSCT were greater (0 ∉ 95% HDI**; Figures 7 and S3**). Most other regions exhibited lower BOLD signals in the second as compared to the first run (WSCT: 197/248, visuomotor: 176/248; **Figure 7**). This included task-activated regions (WSCT: 120, visuomotor: 47), and task-deactivated regions (WSCT: 77, visuomotor: 129), indicating an increase in deactivation. However, the 95% HDI for these comparisons included zero and the magnitude of the between run effect was generally smaller than the overall task effect.

**Figure 7.**
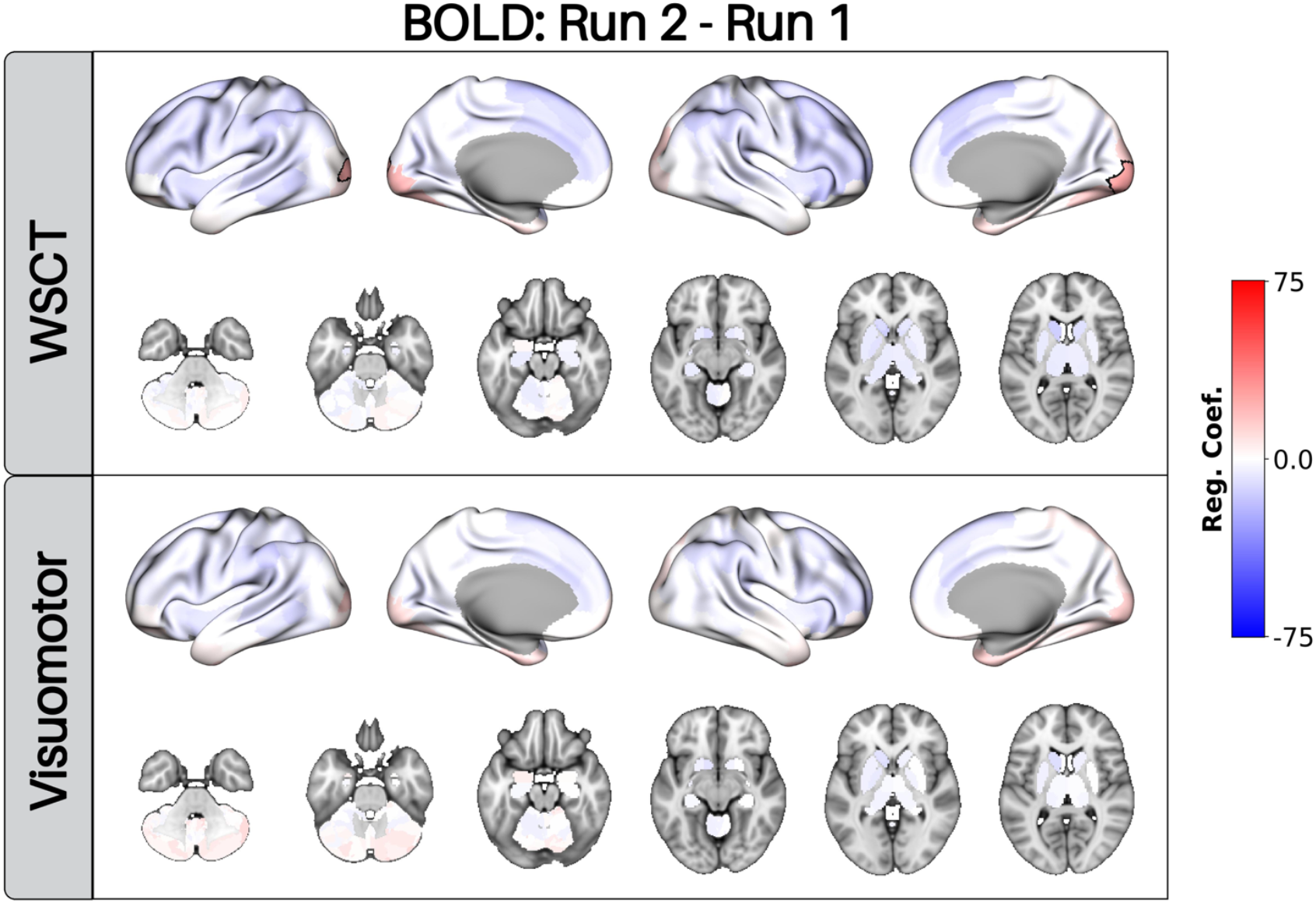
Prolonged task-performance has a minimal impact on fMRI BOLD. Differences between the second and first BOLD runs are displayed for each task. Most regions showed less of a BOLD response in the second run, although the 95% HDI always overlapped 0. There was one parcel in the right visual cortex where the BOLD response to the WSCT was greater during the second run at the 95% HDI level (black outlines). The same color scale is used as **Figure 2** to highlight that run effects are generally much smaller than the main task effects.

## Discussion

Interpreting task-evoked decreases in CBF and BOLD fMRI is challenging because the underlying physiology remains unclear. Using a traditional single bolus FDG PET approach, we found that BOLD deactivations were not accompanied by decreases in CMRglc. Instead, there was a global trend toward elevated CMRglc, with several task-deactivated regions showing significant increases. The lack of task-evoked decreases in CMRglc was consistent across tasks, indicating that task-negative responses are not necessarily driven by reduced metabolic demand.

One proposed explanation for task-negative responses is the vascular steal hypothesis, which posits that deactivations arise from reduced CBF due to activation in nearby regions. Although sensory stimulation can reduce blood flow in areas adjacent to activations (Woolsey et al., 1996; Shmuel et al., 2002), task-negative responses are not always adjacent to activated regions (Shulman et al., 1997). Another hypothesis is that deactivations reflect a reduction in neuronal activity. Most, but not all (Devor et al., 2008), electrophysiology studies report reduced local field potentials or multiunit activity in task-deactivated regions (Shmuel et al., 2006; Lachaux et al., 2008; Hayden et al., 2009; Popa et al., 2009; Jerbi et al., 2010). While extracellular potentials are not always a direct proxy for spiking (Buzsáki et al., 2012), single-unit recordings in task-deactivated regions have shown that decreased fMRI BOLD or extracellular potentials can be accompanied by a decrease in neuronal firing (Hayden et al., 2009; Laurent et al., 2025). However, this is not a universal finding, as a separate study reported increased neuronal firing in task-deactivated regions (Popa et al., 2009).

Recent studies have also highlighted discrepancies between task-evoked changes in BOLD signals and cerebral metabolism. For example, Epp et al. reported that BOLD deactivations can occur without a decrease in cerebral oxygen consumption (Epp et al., 2025). Similarly, in a series of studies using fPET, an infusion-based FDG PET technique, Hahn and colleagues identified task specific dissociations between CMRglc and BOLD signal in the DMN (Godbersen et al., 2023). Some tasks, such as visual stimulation and playing modified Tetris, decreased CMRglc in the DMN (Hahn et al., 2016; Godbersen et al., 2023), whereas a working memory task increased CMRglc (Stiernman et al., 2021).

We did not observe a task-dependent relationship between CMRglc and task-deactived BOLD signals. Rather, widespread increases in CMRglc were observed in both tasks. Differences in approach likely contribute to this discrepancy. fPET is typically analyzed using a generalized linear model, which requires a “baseline” regressor to isolate task-evoked changes in CMRglc (Villien et al., 2014). However, there is evidence that errors in the baseline regressor can induce small (~5%) deactivations (Coursey et al., 2024). In contrast, the bolus approach assumes CMRglc is constant across the scan, an assumption not met here since each task occupied only half of the 60-minute acquisition. As a result, our estimates reflect a mix of rest and task, though the task portion dominates because about two-thirds of the total FDG phosphorylation in the 1-hour scan occurs during the first 30 minutes (Phelps et al., 1979).

Perhaps a more relevant difference between the bolus approach and fPET is how rest and task conditions are defined. fPET acquires both conditions within a single session, eliminating between-session variance, whereas the bolus approach usually requires separate scans on different days. Although sensitivity to between-session variability can be a disadvantage, as some of the variance is likely not physiological, it may also reveal genuine metabolic changes. The widespread increases in CMRglc we observed could indicate an increase in arousal, from a rest scan with no external demands to a task scan requiring sustained alertness and engagement. Although we cannot directly test the role of arousal, as no physiological measures were collected, Madsen et al. reported that task engagement increased whole-brain CMRglc along with heart rate and blood pressure (Madsen et al., 1995). More broadly, our findings align with evidence that arousal induces widespread changes in brain activity (Stringer et al., 2019; Raut et al., 2021) and the observation of widely distributed BOLD fMRI transients at task block onsets and offsets (Fox et al., 2005).

Global activity has often been treated as a nuisance variable in neuroimaging and removed to isolate task-specific effects (Friston et al., 1990). While this is reasonable when the focus is on relative regional effects, it can obscure the broader picture when global activity changes (Borghammer et al., 2008). This is evident in our whole-brain–normalized FDG uptake maps, which showed apparent regional decreases in glucose consumption that are better interpreted as regions where metabolism increased less than the whole-brain average. The removal of global variance may explain why normalized FDG images more closely resembled BOLD fMRI data than quantitative CMRglc maps, particularly with respect to DMN deactivations. In our BOLD fMRI experiment, the baseline was defined by brief (~3 s) inter-trial rest periods, so any state differences across sessions was shared by both task and rest trials. Normalized FDG images may similarly minimize state-related effects, thereby highlighting other processes that contribute to task-evoked changes in glucose metabolism.

While FDG PET reflects the cumulative processes contributing to glucose metabolism, functional magnetic resonance spectroscopy (fMRS) can measure specific metabolites, such as glutamate/glutamine and GABA, in task-deactivated regions (Koush et al., 2022). Two recent fMRS studies reported decreased glutamate concentration in primary visual cortex following stimulation with a small-flickering checkboard, a stimulus that reliably produces a negative BOLD response (Martínez-Maestro et al., 2019; Boillat et al., 2020). In contrast, fMRS studies targeting the DMN have yielded less consistent results; one reported an increase in glutamate/glutamine (Huang et al., 2015), whereas another found increased GABA without a significant change in glutamate/glutamine (Koush et al., 2021). Elevated GABA in task-deactivated regions is consistent with reports of a negative correlation between basal GABA levels and task-evoked BOLD, supporting the idea that inhibitory activity contributes to negative BOLD responses (Northoff et al., 2007).

Increased GABAergic inhibition could also explain elevated CMRglc during tasks. Work in animal models suggests that the rise in glucose consumption during sensory stimulation may occur more so at synapses rather than cell bodies (Schwartz et al., 1979; Kadekaro et al., 1985). One hypothesis, therefore, is that task-negative increases in CMRglc reflect enhanced synaptic GABAergic activity, while decreases in BOLD, CBF, and extracellular electric potentials reflect the resultant reduced glutamatergic activity. However, it is not clear if presynaptic inhibitory activity alone could account for measurable increases in CMRglc as GABAergic activity accounts for only ~20% of glucose oxidation (Patel et al., 2005) and postsynaptic processes are thought to be more energetically costly (Howarth et al., 2012).

Counterintuitively, disinhibition of the task-negative areas could also explain the discrepant change between FDG uptake versus the BOLD signal. Disinhibition of neurons in the task-negative regions would be expected to increase the firing rate, thereby accounting for an increase in CMRglc. However, such an increase in the firing rate does not necessarily imply an increase in local field potentials, which can even decrease despite the higher firing rate due to a loss of synchrony or neuromodulatory changes, and thus manifest as a decrease in the BOLD signal (Popa et al., 2009). Imaging methods capable of resolving glucose metabolism at the cellular level (Zhang et al., 2019) will be necessary to adjudicate between these various hypotheses.

Several limitations should be considered when interpreting our findings. First, although BOLD-negative regions were more likely to show increased CMRglc, small (−5%) decreases could not be ruled out in some areas. As fPET studies have reported small effect sizes in task-negative regions (Godbersen et al., 2023), future work with improved methods is needed for more precise estimates. Another limitation is our use of image-derived input functions. We chose image-derived input functions because acquiring arterial sampling on three separate visits would have placed an unfeasible burden on participants. While increasingly accepted for FDG, image-derived input functions remain susceptible to artifacts owing to partial volume effects and motion (Volpi et al., 2023). Our extraction methods were based on techniques validated against arterial sampling, but differed in key respects (e.g., no venous samples). Although two extraction methods yielded similar results, optimization and standardization of image-derived arterial input functions will be important. Finally, to accommodate simultaneous PET/MR, participants performed both tasks for ~30 minutes, considerably longer than is typical for fPET or fMRI studies. Although prolonged task performance can change the metabolic response to sensory stimulation (Mintun et al., 2002; Vlassenko et al., 2006), we did not find robust changes in the BOLD response between the start and end of the task period, suggesting that any changes in metabolism were minimal.

With these limitations in mind, we cautiously conclude that prolonged task performance does not lead to substantial decreases in CMRglc in regions showing a task-negative BOLD response. Rather, CMRglc actually appears to increase in some task-negative regions as part of a widespread increase in metabolism during task engagement. Whether this widespread increase reflects a brain-wide increase in neuronal activity associated with arousal or another underlying process is an important question for future research.

## Acknowledgements

Funding for this research was provided by the National Institutes of Health/National Institute on Aging grants RF1AG074992 (AGV, MSG) and P50AG0005681 (John C Morris), P01AG026276 (JCM), and P01AG003991 (JCM, Tamme LS Benzinger).

We greatly appreciate Tony Durbin, Jennifer Byers, Kim Casey, Nicholas Metcalf, and Hussain Jafri for their ongoing efforts in participant recruitment and data collection. We are particularly grateful for our research participants and their families for their altruism. We also acknowledge the directors and staff of the Neuroimaging Labs Research Center, Knight Alzheimer’s Disease Research Center, Center for Clinical Imaging Research and the Washington University cyclotron facility for making this research possible.

---

**Figure S1:**
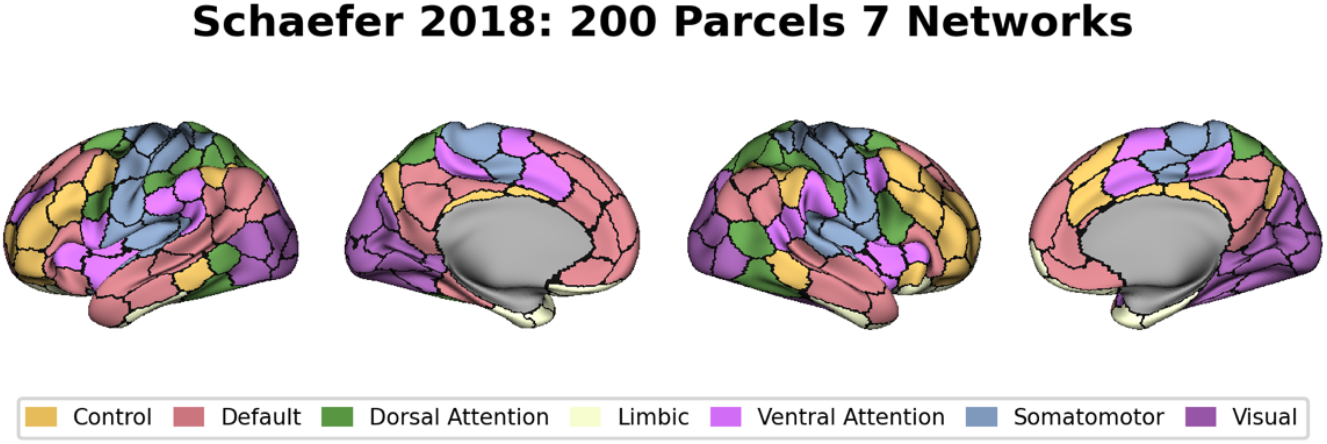
Network assignment of each of the 200 Schaefer parcels.

**Figure S2.**
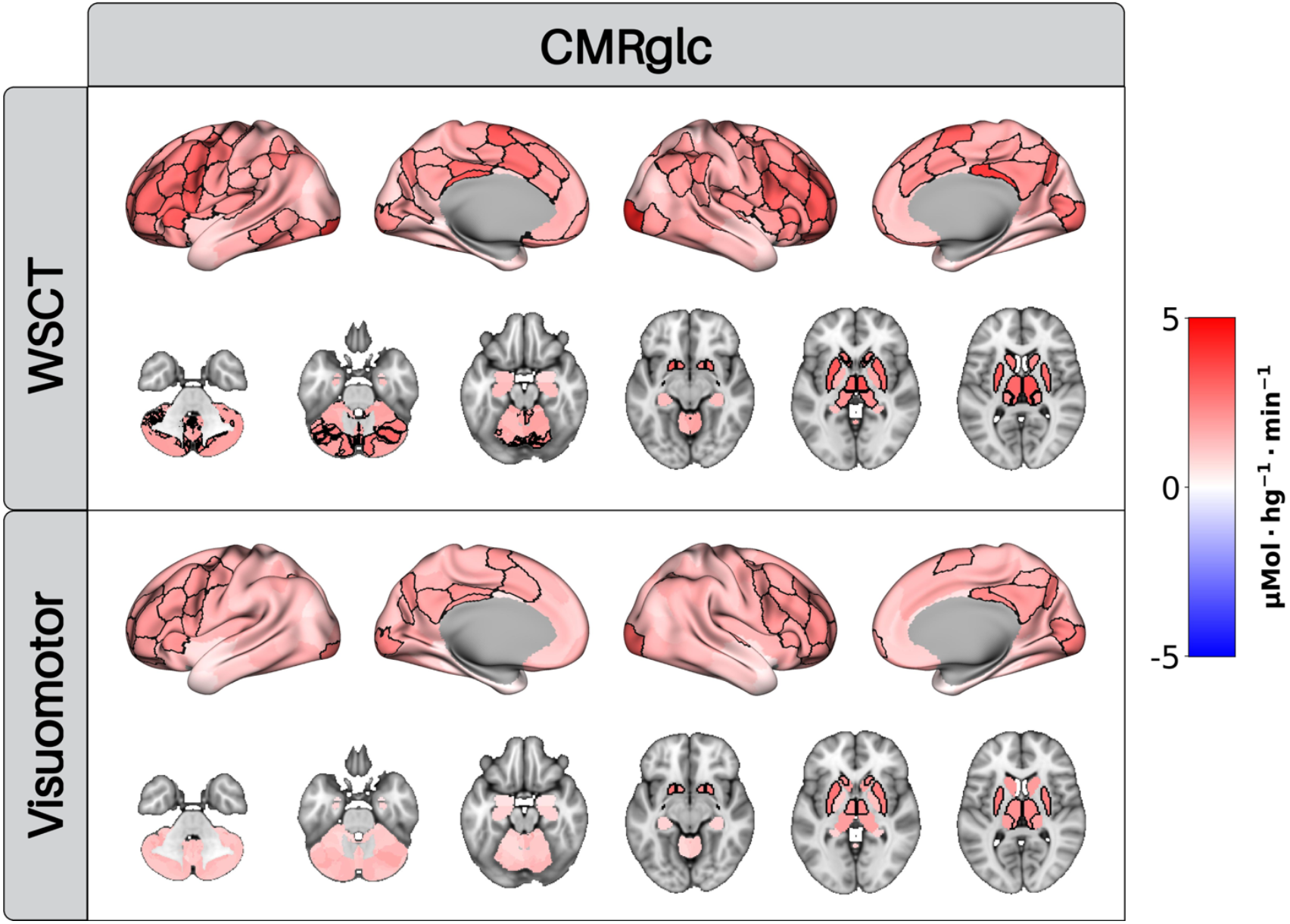
Lack of task-evoked decreases in CMRglc is robust to input function method. Same analysis as in **Figure 2** using an alternative method to extract the image-derived arterial input function (Sari et al., 2017). The results largely mirror those in **Figure 2**, with widespread increases in CMRglc observed in both tasks. There are differences between methods however: more regions met the 95%, particularly for the visuomotor task with this method, and the effect sizes differed, with larger effects for the visuomotor task and smaller effects for the WSCT.

**Figure S3:**
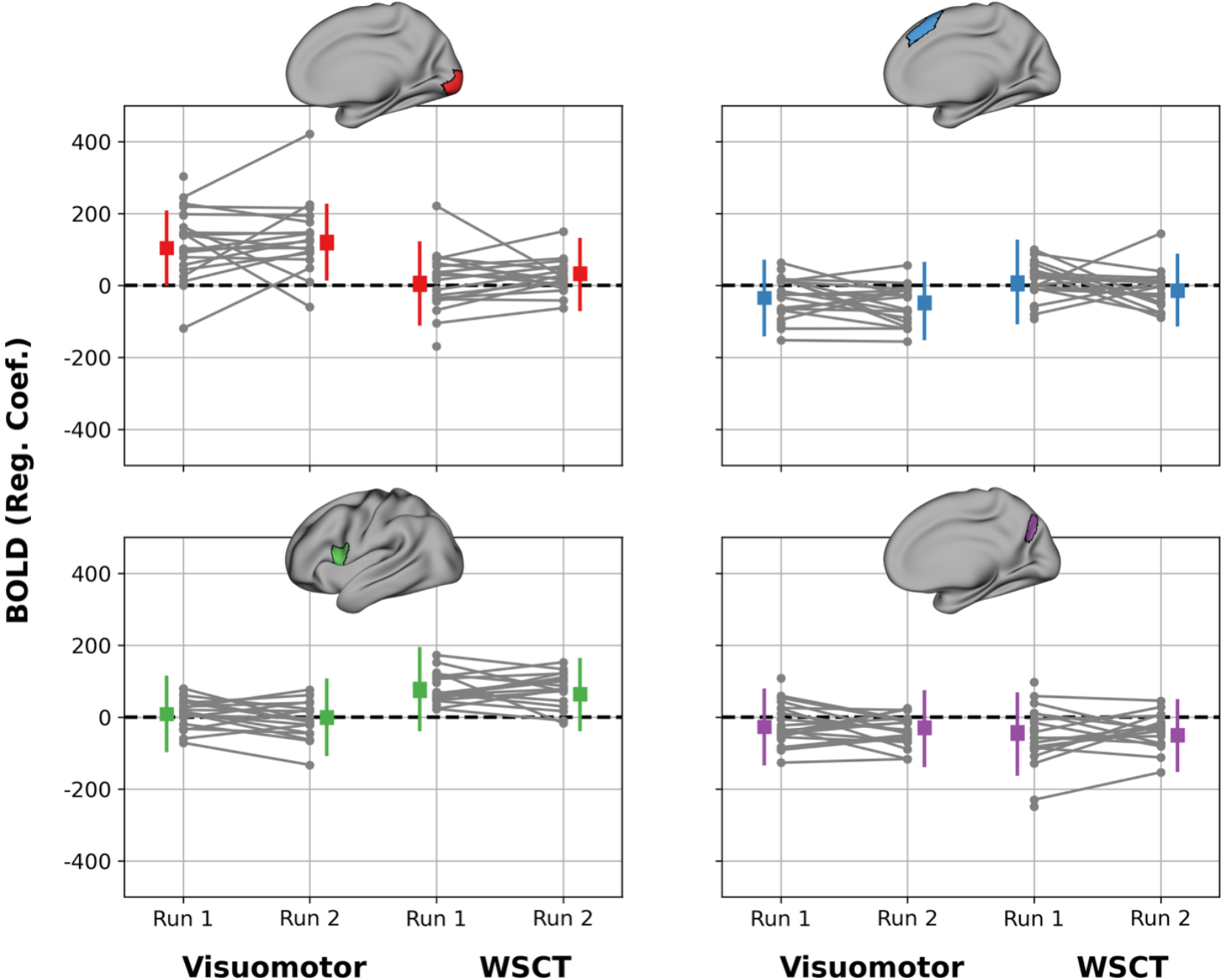
Estimate of task-evoked BOLD activity in four regions of interest. Each gray dot is from single BOLD run, with gray lines connecting data from individual participants. The thick colored lines are the 95% highest density interval (HDI) for each run, while the thin colored lines are the 95% posterior predictive interval. The only region where the HDI excluded zero between runs was the right medial visual cortex (top left), where the second run had a larger BOLD response than the first run during the WSCT.

## Notes

**Conflict of Interest Statement:** AZS is a consultant for Sora Neuroscience LLD.

http://doi.org/10.5281/zenodo.20073537

### Competing Interest Statement

AZS is a consultant for Sora Neuroscience LLD

## References

An H, Lin W (2003) Impact of intravascular signal on quantitative measures of cerebral oxygen extraction and blood volume under normo-and hypercapnic conditions using an asymmetric spin echo approach. Magn Reson Med 50:708–716 Available at: http://www.ncbi.nlm.nih.gov/pubmed/14523956.

Andreasen NC, O’Leary DS, Cizadlo T, Arndt S, Rezai K, Watkins GL, Ponto LL, Hichwa RD (1995) Remembering the past: two facets of episodic memory explored with positron emission tomography. Am J Psychiatry 152:1576–1585 Available at: http://www.ncbi.nlm.nih.gov/pubmed/7485619.

Avants BB, Epstein CL, Grossman M, Gee JC (2008) Symmetric diffeomorphic image registration with cross-correlation: Evaluating automated labeling of elderly and neurodegenerative brain. Med Image Anal 12:26–41.

Avants BB, Tustison NJ, Song G, Cook PA, Klein A, Gee JC (2011) A reproducible evaluation of ANTs similarity metric performance in brain image registration. Neuroimage 54:2033–2044 Available at: 10.1016/j.neuroimage.2010.09.025.

Barch DM, Sabb FW, Carter CS, Braver TS, Noll DC, Cohen JD (1999) Overt verbal responding during fMRI scanning: empirical investigations of problems and potential solutions. Neuroimage 10:642–657 Available at: http://www.ncbi.nlm.nih.gov/pubmed/10600410.

Barnard J, Mcculloch R, Meng X-L (2000) Modeling covariance matrices in terms of standard deviations and correlations, with application to shrinkage. Stat Sin 10:1281–1311.

Birn RM, Bandettini PA, Cox RW, Jesmanowicz A, Shaker R (1998) Magnetic field changes in the human brain due to swallowing or speaking. Magn Reson Med 40:55–60 Available at: http://www.ncbi.nlm.nih.gov/pubmed/9660553.

Blazey T, Snyder AZ, Goyal MS, Vlassenko AG, Raichle ME (2018) A systematic meta-analysis of oxygen-to-glucose and oxygen-to-carbohydrate ratios in the resting human brain. PLoS One 13:e0204242 Available at: http://www.ncbi.nlm.nih.gov/pubmed/30248124.

Boillat Y, Xin L, van der Zwaag W, Gruetter R (2020) Metabolite concentration changes associated with positive and negative BOLD responses in the human visual cortex: A functional MRS study at 7 Tesla. Journal of Cerebral Blood Flow and Metabolism 40:488–500.

Borghammer P, Jonsdottir KY, Cumming P, Ostergaard K, Vang K, Ashkanian M, Vafaee M, Iversen P, Gjedde A (2008) Normalization in PET group comparison studies--the importance of a valid reference region. Neuroimage 40:529–540 Available at: http://www.ncbi.nlm.nih.gov/pubmed/18258457.

Bürkner P-C (2017) brms : An R Package for Bayesian Multilevel Models Using Stan. J Stat Softw 80 Available at: http://www.jstatsoft.org/v80/i01/.

Buzsáki G, Anastassiou CA, Koch C (2012) The origin of extracellular fields and currents--EEG, ECoG, LFP and spikes. Nat Rev Neurosci 13:407–420 Available at: http://www.ncbi.nlm.nih.gov/pubmed/22595786.

Chen G, Xiao Y, Taylor PA, Rajendra JK, Riggins T, Geng F, Redcay E, Cox RW (2019) Handling Multiplicity in Neuroimaging Through Bayesian Lenses with Multilevel Modeling. Neuroinformatics 17:515–545 Available at: http://www.ncbi.nlm.nih.gov/pubmed/30649677.

Coursey SE, Mandeville J, Reed MB, Hartung GA, Garimella A, Sari H, Lanzenberger R, Price JC, Polimeni JR, Greve DN, Hahn A, Chen JE (2024) On the analysis of functional PET (fPET)-FDG: baseline mischaracterization can introduce artifactual metabolic (de)activations. bioRxiv Available at: http://www.ncbi.nlm.nih.gov/pubmed/39484579.

Dai W, Garcia D, de Bazelaire C, Alsop DC (2008) Continuous flow-driven inversion for arterial spin labeling using pulsed radio frequency and gradient fields. Magn Reson Med 60:1488–1497 Available at: http://www.ncbi.nlm.nih.gov/pubmed/19025913.

Dai X, Chen Z, Tian J (2011) Performance evaluation of kinetic parameter estimation methods in dynamic FDG-PET studies. Nucl Med Commun 32:4–16 Available at: http://www.ncbi.nlm.nih.gov/pubmed/21166088.

Dale AM, Fischl B, Sereno MI (1999) Cortical Surface-Based Analysis: I. Segmentation and Surface Reconstruction. Neuroimage 9:179–194 Available at: https://www.sciencedirect.com/science/article/pii/S1053811998903950.

De Francisci M, Silvestri E, Bettinelli A, Volpi T, Goyal MS, Vlassenko AG, Cecchin D, Bertoldo A (2024) EMATA: a toolbox for the automatic extraction and modeling of arterial inputs for tracer kinetic analysis in [18F]FDG brain studies. EJNMMI Phys 11:105 Available at: http://www.ncbi.nlm.nih.gov/pubmed/39715888.

Devor A, Hillman EMC, Tian P, Waeber C, Teng IC, Ruvinskaya L, Shalinsky MH, Zhu H, Haslinger RH, Narayanan SN, Ulbert I, Dunn AK, Lo EH, Rosen BR, Dale AM, Kleinfeld D, Boas DA (2008) Stimulus-induced changes in blood flow and 2-deoxyglucose uptake dissociate in ipsilateral somatosensory cortex. Journal of Neuroscience 28:14347–14357.

Dixon WT (1984) Simple proton spectroscopic imaging. Radiology 153:189–194 Available at: http://www.ncbi.nlm.nih.gov/pubmed/6089263.

Epp SM, Castrillón G, Yuan B, Andrews-Hanna J, Preibisch C, Riedl V (2025) BOLD signal changes can oppose oxygen metabolism across the human cortex. Nat Neurosci Available at: http://www.ncbi.nlm.nih.gov/pubmed/41402521.

Esteban O, Markiewicz CJ, Blair RW, Moodie CA, Isik AI, Erramuzpe A, Kent JD, Goncalves M, DuPre E, Snyder M, Oya H, Ghosh SS, Wright J, Durnez J, Poldrack RA, Gorgolewski KJ (2019) fMRIPrep: a robust preprocessing pipeline for functional MRI. Nat Methods 16:111–116 Available at: http://www.ncbi.nlm.nih.gov/pubmed/30532080.

Fan AP, Jahanian H, Holdsworth SJ, Zaharchuk G (2016) Comparison of cerebral blood flow measurement with [15O]-water positron emission tomography and arterial spin labeling magnetic resonance imaging: A systematic review. J Cereb Blood Flow Metab 36:842–861 Available at: http://www.ncbi.nlm.nih.gov/pubmed/26945019.

Fischl B (2012) FreeSurfer. Neuroimage 62:774–781.

Floberg JM, Mistretta CA, Weichert JP, Hall LT, Holden JE, Christian BT (2012) Improved kinetic analysis of dynamic PET data with optimized HYPR-LR. Med Phys 39:3319–3331.

Fox MD, Snyder AZ, Barch DM, Gusnard DA, Raichle ME (2005) Transient BOLD responses at block transitions. Neuroimage 28:956–966 Available at: http://www.ncbi.nlm.nih.gov/pubmed/16043368.

Friston KJ, Frith CD, Liddle PF, Dolan RJ, Lammertsma AA, Frackowiak RSJ (1990) The relationship between global and local changes in PET scans. Journal of Cerebral Blood Flow and Metabolism 10:458–466.

Frith CD, Friston KJ, Liddle PF, Frackowiak RS (1991) A PET study of word finding. Neuropsychologia 29:1137–1148 Available at: http://www.ncbi.nlm.nih.gov/pubmed/1791928.

Gelman A, Carlin JB, Stern HS, Dunson DB, Vehtari A, Rubin DB (2013) Bayesian Data Analysis. Chapman and Hall/CRC. Available at: https://www.taylorfrancis.com/books/9781439898208.

Godbersen GM, Klug S, Wadsak W, Pichler V, Raitanen J, Rieckmann A, Stiernman L, Cocchi L, Breakspear M, Hacker M, Lanzenberger R, Hahn A (2023) Task-evoked metabolic demands of the posteromedial default mode network are shaped by dorsal attention and frontoparietal control networks.

Gorgolewski K, Burns CD, Madison C, Clark D, Halchenko YO, Waskom ML, Ghosh SS (2011) Nipype: A flexible, lightweight and extensible neuroimaging data processing framework in Python. Front Neuroinform 5.

Gou Y, Golden WC, Lin Z, Shepard J, Tekes A, Hu Z, Li X, Oishi K, Albert M, Lu H, Liu P, Jiang D (2024) Automatic Rejection based on Tissue Signal (ARTS) for motion-corrected quantification of cerebral venous oxygenation in neonates and older adults. Magn Reson Imaging 105:92–99 Available at: http://www.ncbi.nlm.nih.gov/pubmed/37939974.

Greicius MD, Menon V (2004) Default-mode activity during a passive sensory task: uncoupled from deactivation but impacting activation. J Cogn Neurosci 16:1484–1492 Available at: http://www.ncbi.nlm.nih.gov/pubmed/15601513.

Greve DN, Fischl B (2009) NeuroImage Accurate and robust brain image alignment using boundary-based registration. Neuroimage 48:63–72 Available at: 10.1016/j.neuroimage.2009.06.060.

Hahn A, Gryglewski G, Nics L, Hienert M, Rischka L, Vraka C, Sigurdardottir H, Vanicek T, James GM, Seiger R, Kautzky A, Silberbauer L, Wadsak W, Mitterhauser M, Hacker M, Kasper S, Lanzenberger R (2016) Quantification of Task-Specific Glucose Metabolism with Constant Infusion of 18F-FDG. J Nucl Med 57:1933–1940 Available at: http://jnm.snmjournals.org/lookup/doi/10.2967/jnumed.116.176156.

Hahn A, Gryglewski G, Nics L, Rischka L, Ganger S, Sigurdardottir H, Vraka C, Silberbauer L, Vanicek T, Kautzky A, Wadsak W, Mitterhauser M, Hartenbach M, Hacker M, Kasper S, Lanzenberger R (2018) Task-relevant brain networks identified with simultaneous PET/MR imaging of metabolism and connectivity. Brain Struct Funct 223:1369–1378.

Harris CR et al. (2020) Array programming with NumPy. Nature 585:357–362 Available at: 10.1038/s41586-020-2649-2.

Haxby J V, Horwitz B, Ungerleider LG, Maisog JM, Pietrini P, Grady CL (1994) The functional organization of human extrastriate cortex: a PET-rCBF study of selective attention to faces and locations. J Neurosci 14:6336–6353 Available at: http://www.ncbi.nlm.nih.gov/pubmed/7965040.

Hayden BY, Smith D V, Platt ML (2009) Electrophysiological correlates of default-mode processing in macaque posterior cingulate cortex. Proc Natl Acad Sci U S A 106:5948–5953 Available at: http://www.ncbi.nlm.nih.gov/pubmed/19293382.

Henriksen OM, Gjedde A, Vang K, Law I, Aanerud J, Rostrup E (2021) Regional and interindividual relationships between cerebral perfusion and oxygen metabolism. J Appl Physiol (1985) 130:1836–1847 Available at: http://www.ncbi.nlm.nih.gov/pubmed/33830816.

Hoopes A, Mora JS, Dalca A V, Fischl B, Hoffmann M (2022) SynthStrip: skull-stripping for any brain image. Neuroimage 260:119474 Available at: http://www.ncbi.nlm.nih.gov/pubmed/35842095.

Howarth C, Gleeson P, Attwell D (2012) Updated energy budgets for neural computation in the neocortex and cerebellum. J Cereb Blood Flow Metab 32:1222–1232 Available at: http://www.ncbi.nlm.nih.gov/pubmed/22434069.

Huang Z, Davis HH, Yue Q, Wiebking C, Duncan NW, Zhang J, Wagner N-F, Wolff A, Northoff G (2015) Increase in glutamate/glutamine concentration in the medial prefrontal cortex during mental imagery: A combined functional mrs and fMRI study. Hum Brain Mapp 36:3204–3212 Available at: http://www.ncbi.nlm.nih.gov/pubmed/26059006.

Jenkinson M (2003) Fast, automated, N-dimensional phase-unwrapping algorithm. Magn Reson Med 49:193–197.

Jenkinson M, Bannister P, Brady M, Smith S (2002) Improved optimization for the robust and accurate linear registration and motion correction of brain images. Neuroimage 17:825–841 Available at: http://www.ncbi.nlm.nih.gov/pubmed/12377157.

Jenkinson M, Beckmann CF, Behrens TEJ, Woolrich MW, Smith SM (2012) FSL. Neuroimage 62:782–790 Available at: https://linkinghub.elsevier.com/retrieve/pii/S1053811911010603.

Jenkinson M, Smith S (2001) A global optimisation method for robust affine registration of brain images. Med Image Anal 5:143–156 Available at: https://www.sciencedirect.com/science/article/pii/S1361841501000366.

Jerbi K, Vidal JR, Ossandon T, Dalal SS, Jung J, Hoffmann D, Minotti L, Bertrand O, Kahane P, Lachaux J-P (2010) Exploring the electrophysiological correlates of the default-mode network with intracerebral EEG. Front Syst Neurosci 4:27 Available at: http://www.ncbi.nlm.nih.gov/pubmed/20661461.

Jiang D, Deng S, Franklin CG, O’Boyle M, Zhang W, Heyl BL, Pan L, Jerabek PA, Fox PT, Lu H (2021) Validation of T2-based oxygen extraction fraction measurement with 15 O positron emission tomography. Magn Reson Med 85:290–297 Available at: http://www.ncbi.nlm.nih.gov/pubmed/32643207.

Jiang D, Lu H (2022) Cerebral oxygen extraction fraction MRI: Techniques and applications. Magn Reson Med 88:575–600 Available at: http://www.ncbi.nlm.nih.gov/pubmed/35510696.

Jung J, Jerbi K, Ossandon T, Ryvlin P, Isnard J, Bertrand O, Guénot M, Mauguière F, Lachaux J-P (2010) Brain responses to success and failure: Direct recordings from human cerebral cortex. Hum Brain Mapp 31:1217–1232 Available at: http://www.ncbi.nlm.nih.gov/pubmed/20120013.

Kadekaro M, Crane AM, Sokoloff L (1985) Differential effects of electrical stimulation of sciatic nerve on metabolic activity in spinal cord and dorsal root ganglion in the rat. Proc Natl Acad Sci U S A 82:6010–6013 Available at: http://www.ncbi.nlm.nih.gov/pubmed/3862113.

Kay M (2024) ggdist: Visualizations of Distributions and Uncertainty in the Grammar of Graphics. IEEE Trans Vis Comput Graph 30:414–424 Available at: http://www.ncbi.nlm.nih.gov/pubmed/37883271.

Klein A, Ghosh SS, Bao FS, Giard J, Häme Y, Stavsky E, Lee N, Rossa B, Reuter M, Neto EC, Keshavan A (2017) Mindboggling morphometry of human brains. PLoS Comput Biol 13:e1005350 Available at: https://journals.plos.org/ploscompbiol/article?id=10.1371/journal.pcbi.1005350.

Koush Y, de Graaf RA, Kupers R, Dricot L, Ptito M, Behar KL, Rothman DL, Hyder F (2021) Metabolic underpinnings of activated and deactivated cortical areas in human brain. Journal of Cerebral Blood Flow and Metabolism.

Koush Y, Rothman DL, Behar KL, de Graaf RA, Hyder F (2022) Human brain functional MRS reveals interplay of metabolites implicated in neurotransmission and neuroenergetics. Journal of Cerebral Blood Flow and Metabolism 42:911–934.

Lachaux JP, Jung J, Mainy N, Dreher JC, Bertrand O, Baciu M, Minotti L, Hoffmann D, Kahane P (2008) Silence is golden: transient neural deactivation in the prefrontal cortex during attentive reading. Cereb Cortex 18:443–450 Available at: http://www.ncbi.nlm.nih.gov/pubmed/17617656.

Laurent M-A, Jacques C, Yan X, Jurczynski P, Colnat-Coulbois S, Maillard L, Le Cam S, Ranta R, Cottereau BR, Koessler L, Jonas J, Rossion B (2025) A tight relationship between BOLD fMRI activation/deactivation and increase/decrease in single neuron responses in human association cortex. Elife 14 Available at: http://www.ncbi.nlm.nih.gov/pubmed/40793082.

Lewandowski D, Kurowicka D, Joe H (2009) Generating random correlation matrices based on vines and extended onion method. J Multivar Anal 100:1989–2001.

Logothetis NK, Pauls J, Augath M, Trinath T, Oeltermann A (2001) Neurophysiological investigation of the basis of the fMRI signal. Nature 412:150–157 Available at: http://www.ncbi.nlm.nih.gov/pubmed/11449264.

Lu H, Ge Y (2008) Quantitative evaluation of oxygenation in venous vessels using T2-Relaxation-Under-Spin-Tagging MRI. Magn Reson Med 60:357–363 Available at: http://www.ncbi.nlm.nih.gov/pubmed/18666116.

Lu H, Xu F, Grgac K, Liu P, Qin Q, Van Zijl P (2012) Calibration and validation of TRUST MRI for the estimation of cerebral blood oxygenation. Magn Reson Med 67:42–49.

Madsen PL, Hasselbalch SG, Hagemann LP, Olsen KS, Bülow J, Holm S, Wildschiødtz G, Paulson OB, Lassen NA (1995) Persistent resetting of the cerebral oxygen/glucose uptake ratio by brain activation: Evidence obtained with the Kety-Schmidt technique. Journal of Cerebral Blood Flow and Metabolism 15:485–491.

Makowski D, Ben-Shachar MS, Chen SHA, Lüdecke D (2019) Indices of Effect Existence and Significance in the Bayesian Framework. Front Psychol 10:2767 Available at: http://www.ncbi.nlm.nih.gov/pubmed/31920819.

Martínez-Maestro M, Labadie C, Möller HE (2019) Dynamic metabolic changes in human visual cortex in regions with positive and negative blood oxygenation level-dependent response. Journal of Cerebral Blood Flow and Metabolism 39:2295–2307.

McKiernan KA, Kaufman JN, Kucera-Thompson J, Binder JR (2003) A parametric manipulation of factors affecting task-induced deactivation in functional neuroimaging. J Cogn Neurosci 15:394–408 Available at: http://www.ncbi.nlm.nih.gov/pubmed/12729491.

Mintun MA, Vlassenko AG, Shulman GL, Snyder AZ (2002) Time-related increase of oxygen utilization in continuously activated human visual cortex. Neuroimage 16:531–537.

Nettekoven C, Zhi D, Shahshahani L, Pinho AL, Saadon-Grosman N, Buckner RL, Diedrichsen J (2024) A hierarchical atlas of the human cerebellum for functional precision mapping. Nat Commun 15:8376 Available at: http://www.ncbi.nlm.nih.gov/pubmed/39333089.

Northoff G, Walter M, Schulte RF, Beck J, Dydak U, Henning A, Boeker H, Grimm S, Boesiger P (2007) GABA concentrations in the human anterior cingulate cortex predict negative BOLD responses in fMRI. Nat Neurosci 10:1515–1517.

Otto Henriksen XM, Vestergaard MB, Lindberg U, Aachmann-Andersen NJ, Lisbjerg K, Christensen SJ, Rasmussen P, Olsen N V, Forman JL W Larsson HB, Law I (2018) Interindividual and regional relationship between cerebral blood flow and glucose metabolism in the resting brain. J Appl Physiol 125:1080–1089 Available at: http://www.jappl.org.

Patel AB, de Graaf RA, Mason GF, Rothman DL, Shulman RG, Behar KL (2005) The contribution of GABA to glutamateglutamine cycling and energy metabolism in the rat cortex in vivo. Available at: www.pnas.orgcgidoi10.1073pnas.0501703102.

Pedregosa F, Varoquaux G, Gramfort A, Michel V, Thirion B, Grisel O, Blondel M, Prettenhofer P, Weiss R, Dubourg V, Vanderplas J, Passos A, Cournapeau D, Brucher M, Perrot M, Duchesnay É (2011) Scikit-Learn: Machine Learning in Python. J Mach Learn Res 12:2825– 2830.

Peirce J, Gray JR, Simpson S, MacAskill M, Höchenberger R, Sogo H, Kastman E, Lindeløv JK (2019) PsychoPy2: Experiments in behavior made easy. Behav Res Methods 51:195–203.

Phelps ME, Huang SC, Hoffman EJ, Selin C, Sokoloff L, Kuhl DE (1979) Tomographic measurement of local cerebral glucose metabolic rate in humans with (F-18)2-fluoro-2-deoxy-D-glucose: Validation of method. Ann Neurol 6:371–388.

Popa D, Popescu AT, Paré D (2009) Contrasting activity profile of two distributed cortical networks as a function of attentional demands. Journal of Neuroscience 29:1191–1201.

R Core Team (2024) R: A Language and Environment for Statistical Computing. Available at: https://www.R-project.org/.

Raichle ME, MacLeod AM, Snyder AZ, Powers WJ, Gusnard DA, Shulman GL (2001) A default mode of brain function. Proc Natl Acad Sci U S A 98:676–682 Available at: http://www.ncbi.nlm.nih.gov/pubmed/11209064.

Raut R V, Snyder AZ, Mitra A, Yellin D, Fujii N, Malach R, Raichle ME (2021) Global waves synchronize the brain’s functional systems with fluctuating arousal. Sci Adv 7 Available at: http://www.ncbi.nlm.nih.gov/pubmed/34290088.

Reuter M, Rosas HD, Fischl B (2010) Highly accurate inverse consistent registration: A robust approach. Neuroimage 53:1181–1196.

Rischka L, Gryglewski G, Pfaff S, Vanicek T, Hienert M, Klöbl M, Hartenbach M, Haug A, Wadsak W, Mitterhauser M, Hacker M, Kasper S, Lanzenberger R, Hahn A (2018) Reduced task durations in functional PET imaging with [18F]FDG approaching that of functional MRI. Neuroimage 181:323–330.

Rizzo G, Turkheimer FE, Bertoldo A (2013) Multi-scale hierarchical approach for parametric mapping: Assessment on multi-compartmental models. Neuroimage 67:344–353 Available at: 10.1016/j.neuroimage.2012.11.045.

Sari H, Erlandsson K, Law I, Larsson HBW, Ourselin S, Arridge S, Atkinson D, Hutton BF (2017) Estimation of an image derived input function with MR-defined carotid arteries in FDG-PET human studies using a novel partial volume correction method. Journal of Cerebral Blood Flow and Metabolism 37:1398–1409.

Schaefer A, Kong R, Gordon EM, Laumann TO, Zuo X-N, Holmes AJ, Eickhoff SB, Yeo BTT (2018) Local-Global Parcellation of the Human Cerebral Cortex from Intrinsic Functional Connectivity MRI. Cereb Cortex 28:3095–3114 Available at: http://www.ncbi.nlm.nih.gov/pubmed/28981612.

Scholkmann F, Gerber U, Wolf M, Wolf U (2013) End-tidal CO2: an important parameter for a correct interpretation in functional brain studies using speech tasks. Neuroimage 66:71–79 Available at: http://www.ncbi.nlm.nih.gov/pubmed/23099101.

Schwartz WJ, Smith CB, Davidsen L, Savaki H, Sokoloff L, Mata M, Fink DJ, Gainer H (1979) Metabolic mapping of functional activity in the hypothalamo-neurohypophysial system of the rat. Science 205:723–725 Available at: http://www.ncbi.nlm.nih.gov/pubmed/462184.

Shmuel A, Augath M, Oeltermann A, Logothetis NK (2006) Negative functional MRI response correlates with decreases in neuronal activity in monkey visual area V1. Nat Neurosci 9:569–577 Available at: http://www.ncbi.nlm.nih.gov/pubmed/16547508.

Shmuel A, Yacoub E, Pfeuffer J, Van de Moortele PF, Adriany G, Hu X, Ugurbil K (2002) Sustained negative BOLD, blood flow and oxygen consumption response and its coupling to the positive response in the human brain. Neuron 36:1195–1210 Available at: http://www.ncbi.nlm.nih.gov/pubmed/12495632.

Shulman GL, Fiez JA, Corbetta M, Buckner RL, Miezin FM, Raichle ME, Petersen SE (1997) Common Blood Flow Changes across Visual Tasks: II. Decreases in Cerebral Cortex. J Cogn Neurosci 9:648–663 Available at: http://www.ncbi.nlm.nih.gov/pubmed/23965122.

Smith SM (2002) Fast robust automated brain extraction. Hum Brain Mapp 17:143–155.

Sokoloff L (1999) Energetics of functional activation in neural tissues. Neurochem Res 24:321–329 Available at: http://www.ncbi.nlm.nih.gov/pubmed/9972882.

Stan Development Team (2024) RStan: the R interface to Stan. Available at: https://mc-stan.org/.

Stiernman LJ, Grill F, Hahn A, Rischka L, Lanzenberger R, Lundmark VP, Riklund K, Axelsson J, Rieckmann A (2021) Dissociations between glucose metabolism and blood oxygenation in the human default mode network revealed by simultaneous PET-fMRI. Proc Natl Acad Sci U S A 118.

Stringer C, Pachitariu M, Steinmetz N, Reddy CB, Carandini M, Harris KD (2019) Spontaneous behaviors drive multidimensional, brainwide activity. Science 364:255 Available at: http://www.ncbi.nlm.nih.gov/pubmed/31000656.

Tian Y, Margulies DS, Breakspear M, Zalesky A (2020) Topographic organization of the human subcortex unveiled with functional connectivity gradients. Nat Neurosci 23:1421–1432 Available at: http://www.ncbi.nlm.nih.gov/pubmed/32989295.

Villien M, Wey HY, Mandeville JB, Catana C, Polimeni JR, Sander CY, Zürcher NR, Chonde DB, Fowler JS, Rosen BR, Hooker JM (2014) Dynamic functional imaging of brain glucose utilization using fPET-FDG. Neuroimage 100:192–199 Available at: 10.1016/j.neuroimage.2014.06.025.

Virtanen P et al. (2020) SciPy 1.0: fundamental algorithms for scientific computing in Python. Nat Methods 17:261–272.

Vlassenko AG, Rundle MM, Mintun MA (2006) Human brain glucose metabolism may evolve during activation: Findings from a modified FDG PET paradigm. Neuroimage 33:1036–1041.

Volpi T, Maccioni L, Colpo M, Debiasi G, Capotosti A, Ciceri T, Carson RE, DeLorenzo C, Hahn A, Knudsen GM, Lammertsma AA, Price JC, Sossi V, Wang G, Zanotti-Fregonara P, Bertoldo A, Veronese M (2023) An update on the use of image-derived input functions for human PET studies: new hopes or old illusions? EJNMMI Res 13:97 Available at: http://www.ncbi.nlm.nih.gov/pubmed/37947880.

Woolrich MW, Ripley BD, Brady M, Smith SM (2001) Temporal autocorrelation in univariate linear modeling of FMRI data. Neuroimage 14:1370–1386.

Woolsey TA, Rovainen CM, Cox SB, Henegar MH, Liang GE, Liu D, Moskalenko YE, Sui J, Wei L (1996) Neuronal units linked to microvascular modules in cerebral cortex: response elements for imaging the brain. Cereb Cortex 6:647–660 Available at: http://www.ncbi.nlm.nih.gov/pubmed/8921201.

Wu HM, Bergsneider M, Glenn TC, Yeh E, Hovda DA, Phelps ME, Huang SC (2003) Measurement of the global lumped constant for 2-Deoxy-2-[18F]fluoro-D-glucose in normal human brain using [15O]water and 2-deoxy-2-[18F]fluoro-D-glucose positron emission tomography imaging: A method with validation based on Multiple methodologies. Mol Imaging Biol 5:32–41.

Wymer DT, Patel KP, Burke WF, Bhatia VK (2020) Phase-Contrast MRI: Physics, Techniques, and Clinical Applications. Radiographics 40:122–140 Available at: http://www.ncbi.nlm.nih.gov/pubmed/31917664.

Yaqub M, Boellaard R, Kropholler MA, Lammertsma AA (2006) Optimization algorithms and weighting factors for analysis of dynamic PET studies. Phys Med Biol 51:4217–4232 Available at: http://www.ncbi.nlm.nih.gov/pubmed/16912378.

Yushkevich PA, Piven J, Hazlett HC, Smith RG, Ho S, Gee JC, Gerig G (2006) User-guided 3D active contour segmentation of anatomical structures: significantly improved efficiency and reliability. Neuroimage 31:1116–1128 Available at: http://www.ncbi.nlm.nih.gov/pubmed/16545965.

Zhang L, Shi L, Shen Y, Miao Y, Wei M, Qian N, Liu Y, Min W (2019) Spectral tracing of deuterium for imaging glucose metabolism. Nat Biomed Eng 3:402–413 Available at: http://www.ncbi.nlm.nih.gov/pubmed/31036888.

Zhang Y, Brady M, Smith S (2001) Segmentation of brain MR images through a hidden Markov random field model and the expectation-maximization algorithm. IEEE Trans Med Imaging 20:45–57.

